# Nested neural circuits generate distinct acoustic signals during Drosophila courtship

**DOI:** 10.1101/2023.08.30.555537

**Authors:** Joshua L. Lillvis, Kaiyu Wang, Hiroshi M. Shiozaki, Min Xu, David L. Stern, Barry J. Dickson

## Abstract

Many motor control systems generate multiple movements using a common set of muscles. How are premotor circuits able to flexibly generate diverse movement patterns? Here, we characterize the neuronal circuits that drive the distinct courtship songs of *Drosophila melanogaster*. Male flies vibrate their wings towards females to produce two different song modes – pulse and sine song – which signal species identity and male quality. Using cell-type specific genetic reagents and the connectome, we provide a cellular and synaptic map of the circuits in the male ventral nerve cord that generate these songs and examine how activating or inhibiting each cell type within these circuits affects the song. Our data reveal that the song circuit is organized into two nested feed-forward pathways, with extensive reciprocal and feed-back connections. The larger network produces pulse song, the more complex and ancestral song form. A subset of this network produces sine song, the simpler and more recent form. Such nested organization may be a common feature of motor control circuits in which evolution has layered increasing flexibility on to a basic movement pattern.

## Introduction

Many animal behaviors require differential activation of the same set of muscles to produce distinct motor patterns depending on the task at hand. Such motor output is often highly dynamic, responding to moment-to-moment changes in internal drives and environmental cues. This motor flexibility is essential, for example, to travel over varying terrain or to execute skilled hand movements. Another striking example is auditory communication, which many species use to coordinate their social behaviors. Most animals have one primary sound-producing apparatus, and the amount of information that can be transmitted from one individual to another is limited by the number of distinct sounds this apparatus can produce. How is this extraordinary versatility implemented in motor control circuits that act through a common set of muscles? We address this question here in the context of auditory communication during *Drosophila* courtship.

Male flies sing to females during courtship by vibrating one or both wings ^1,2^. *D. melanogaster* males produce two primary song types: pulse song, which consists of trains of discrete ∼220 Hz pulses that recur every 35 milliseconds, and a continuous ∼150 Hz hum called sine song ^1,3,4^ (Figure 1A). Male flies decide whether and how to sing on a moment-to-moment basis depending on internal drives, female behavior, and other environmental cues. For example, previous experience of the male can alter the likelihood of whether the male will sing ^5–7^, and each song type is preferentially produced and modulated in different contexts during courtship ^8–10^. Females evaluate the songs to recognize conspecific males and assess their quality before deciding whether or not to mate ^11–14^. The rich structure of these courtship songs, together with the powerful genetic tools and connectomics resources available in *Drosophila*, make this an attractive model system to investigate how motor control circuits produce distinct sounds.

**Figure 1.**
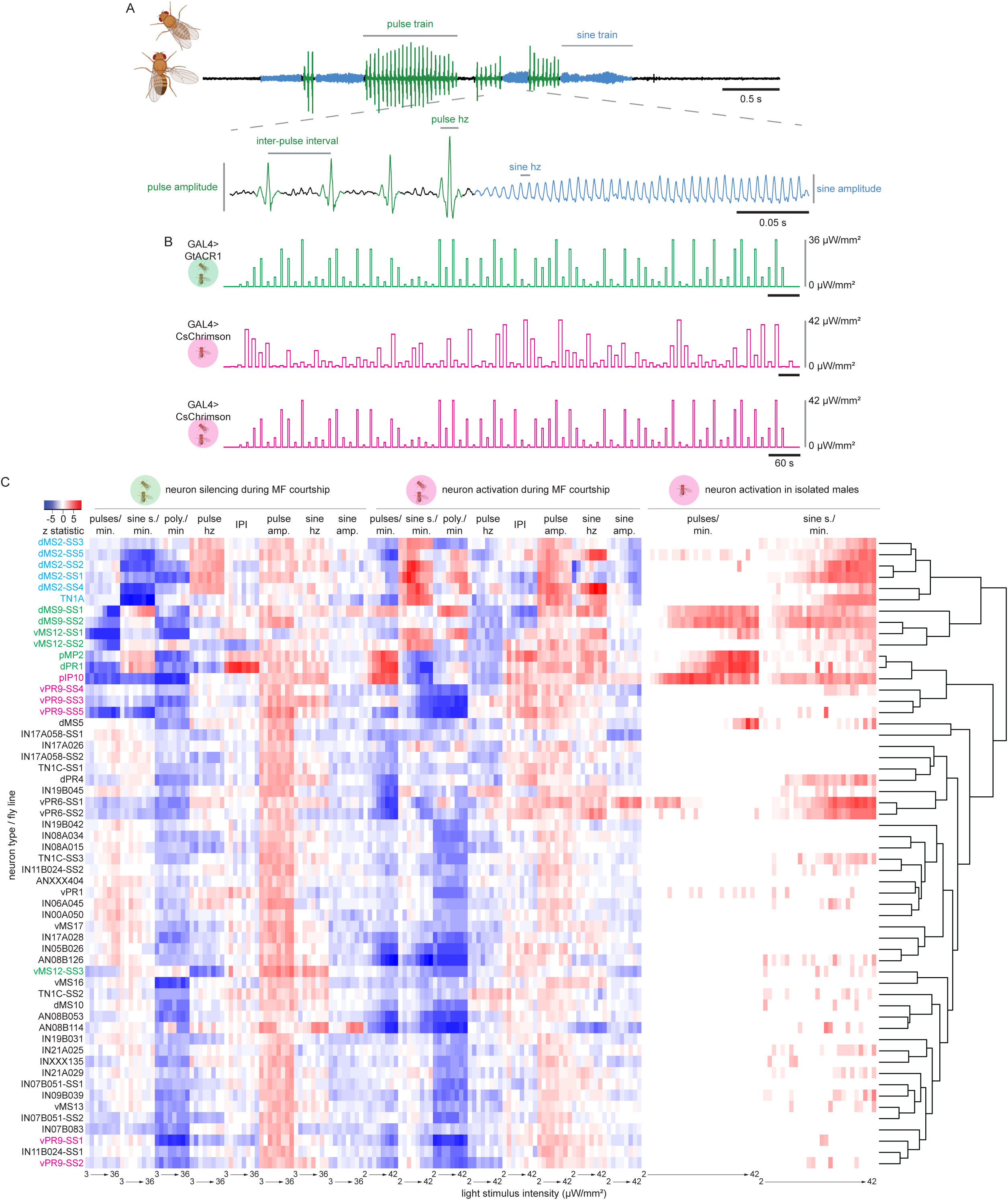
A) Representative example of D. melanogaster courtship song. Males extend and vibrate a single wing while courting females to produce song composed of two primary types: pulse (green) and sine (blue). Pulse trains consist of a series of discreet pulses with a stereotypical frequency, amplitude, and inter-pulse interval. Sine trains consist of a continuous sinusoidal wave with stereotypical frequency and amplitude. B) Behavior assays used to evaluate neuron functions. Single males were paired with single females and neurons were optogenetically silenced using GtACR1. Two different optogenetic CsChrimson activation assays were used: single isolated males and single males paired with single females. For optogenetic assays the timing and intensity of the light stimulation protocols are indicated. GtACR1 was activated by 525 nm green light (green) and CsChrimson was activated by 625 nm red light (magenta). C) Hierarchical clustering of all tested fly lines based on optogenetic experiment song phenotypes. The change in each song characteristic in response to a range of optogenetic silencing and activation light stimuli compared to no stimulus is shown (represented by Dunn’s z test statistic). Each row is a cell-type-specific fly line. Cell types that are required for sine song (cyan), pulse song (green), and both song types (magenta) are indicated.

Courtship behaviors, including song, are controlled largely by a group of approximately 2000 neurons that express either or both of the two sex-determination genes, *fruitless* (*fru*) and *doublesex* (*dsx*) ^15–19^. Genetic driver lines that label specific subsets of these neurons have been instrumental in revealing the functional organization of the circuit. These studies have defined olfactory, gustatory, visual, and auditory pathways that underlie sensorimotor transformations critical to song output ^12,14,28–30,20–27^, as well as key components of the courtship decision making circuitry in the brain ^31–33^. The patterns of muscle and motor neuron activation during pulse and sine song have also been characterized ^34–39^. Still poorly understood, however, are the motor circuits in the ventral nerve cord (VNC, the analog of the vertebrate spinal cord) that transform the descending signals from the brain into the distinct patterns of muscle activation that shape each song modality and feature.

Here we identify many of the VNC neurons that form the *D. melanogaster* song circuit by optogenetically manipulating their activity and quantitatively analyzing the impact of altered neuronal activity on courtship song (Figure 1B). We assess the functions of over 40 distinct cell types by recording and analyzing a total of more than 1800 hours of songs from over 5000 male flies. We then trace the synaptic connections of the identified song neurons in the male VNC connectome (MANC) ^40–42^. These anatomical and functional data define a highly interconnected core song circuit composed of eight *fru* and/or *dsx*-expressing (*fru*^+^/*dsx*^+^) cell types. These neurons are largely organized into two nested feed-forward pathways. The smaller of these pathways generates the simpler sine song; the larger produces the more elaborate pulse song. We also identify small number of additional cell types that may shape specific features of either sine or pulse song or may function in rapid switching between the two song types. We suggest that such a nested circuit architecture may be common theme across animals in the motor circuits that generate diverse movement patterns from a single set of muscles.

## Results

### Identification and functional classification of song neurons in the male ventral nerve cord

A male fly moves its wings in similar ways during flight and courtship song, which are produced by an overlapping set of wing muscles ^39^. However, the descending neurons that drive flight and song make connections to mostly different subsets of interneurons in the male VNC ^41^. To identify neurons in the song circuit, we relied on previous evidence that *fru^+^* and *dsx^+^* neurons in the VNC are required to produce courtship song ^31,35^. We identified neurons within the MANC volume that appeared anatomically similar to *fru*^+^ and *dsx*^+^ neurons in the VNC observed from light microscope surveys (Sup. Table 1) ^31,35,43,44^.

To enable functional analysis of these neurons, we generated a collection of split-GAL4 fly lines that drove expression cleanly in candidate song neurons in the VNC, including 17 of 22 *fru^+^* and *dsx*^+^ cell types identified in the MANC, but also many *fru*^-^ *dsx*^-^ neurons that innervate the wing neuropil. In total, we generated 58 lines covering at least 40 distinct cell types identified in the MANC (Sup. Table 1, Sup. Figure 1-3). We then used these new lines in a series of optogenetic neuronal silencing and activation experiments. We performed three distinct activity perturbation experiments with each line (Figure 1B). To determine whether neuronal activity was necessary for normal song production, we transiently silenced these neurons in open-loop in males that were actively courting a virgin female using the green-light-sensitive inward rectifying chloride channel GtACR1 ^45^. To determine whether neuronal activity was sufficient for song production, we transiently activated these neurons in isolated males using the red-light-sensitive channelrhodopsin CsChrimson ^46^. To determine whether neuronal activation altered ongoing song, we transiently activated these neurons using CsChrimson in males as they courted virgin females. Each of these activity perturbation experiments was performed across a range of stimulus intensities from 0.3 to 42 μW/mm^2^.

We automatically annotated courtship songs in all of these experiments using SongExplorer ^47^, which uses a trained convolutional neural network model to detect courtship song and decompose it into its pulse and sine components. SongExplorer allowed us to quantify the amount of pulse and sine song in each recording, as well as the characteristics of specific song features, including the inter-pulse interval, the number of cycles per pulse, and the carrier frequencies and amplitudes of both pulse and sine song. If a song parameter was perturbed in any of these experiments, it generally varied monotonically with stimulus intensity, although in some cases the strongest phenotype was observed at intermediate stimulus intensities. For those cell types for which we were able to test multiple split-GAL4 driver lines, the shape of dose-response curves was generally consistent across driver lines, but the response at any given stimulus intensity often differed. Quantitative differences between independent driver lines targeting the same cell type may reflect differences in the levels or timing of effector expression, or in some cases differences in the specific subset of neurons labelled within a common cell type. To enable phenotypic comparisons across multiple parameters, cell types and driver lines, we therefore calculated the Dunn’s test z-statistic across all flies and trials for each experimental condition. For each line, we then selected the maximum absolute z-statistic across all stimulus intensities. These values were then used for hierarchical clustering of the entire dataset (Figure 1C).

This analysis defined a core set of eight neuronal cell types that grouped approximately into three functionally related subsets (Figure 1C): two cell types principally involved in sine song production (cyan in Figure 1C), four cell types specifically required for pulse song (green), and two cell types (magenta) with more complex roles in both pulse and sine song. Several additional cell types modulated specific song features without necessarily changing the overall amount of either pulse or sine song. In the following sections, we describe each of the three core song circuit subgroups as well as the neurons that shape specific song features. We then consider these functional data in the light of their neuronal connectivity.

### Neurons that control sine song

Silencing either the TN1A or dMS2 cells significantly reduced the amount of sine song produced by courting males but did not significantly alter the amount of pulse song (Figure 2, Sup. Figure 4). Conversely, activating either cell type in an isolated male was sufficient to elicit sine song but produced little or no pulse song (Figure 2, Sup. Figure 4). Activation in courting males increased the amount of sine song at the expense of pulse song (Sup. Figure 4).

**Figure 2.**
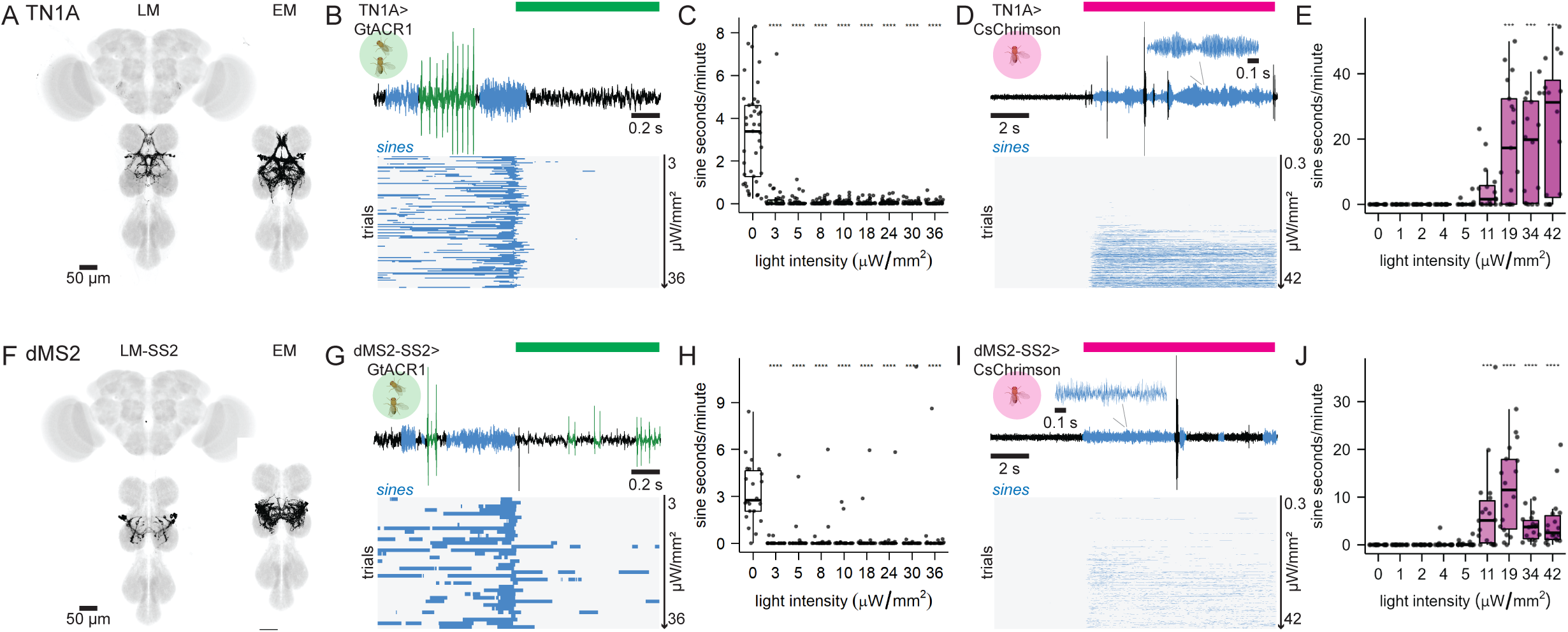
Neuron types necessary for sine song production. A, left) A representative image of TN1A-GAL4 expression. A, right) TN1A in the EM volume. B) Response of courting TN1A>GtACR1 males to green light stimuli that were producing sine at stimulus onset, (top) a representative example, (bottom) ethogram of sine (blue) over time with each row showing a fly/trial and rows sorted by increasing light intensity. C) Quantification of amount of sine in seconds/minute of recording produced by courting TN1A>GtACR1 males without green light (0) and at increasing green light intensities. D) Response of isolated TN1A>CsChrimson males to red light stimuli, (top) a representative example, (bottom) ethogram of sine over time with each row showing a fly/trial and rows sorted by increasing light intensity. E) Quantification of sine seconds/minute produced by isolated TN1A>CsChrimson males without red light (0) and at increasing red light intensities. F-J) Same as A-E), but for dMS2. Pulse and sine trains are indicated in green and blue, respectively. p-values calculated via Kruskal-Wallis tests and Dunn’s pairwise comparisons with Bonferroni correction. *p<0.05, **p<0.005, ***p<0.0005, ****p<0.00005.

TN1A is a heterogeneous group of cholinergic *fru*^+^ and *dsx*^+^ neurons, together forming one subset of the larger TN1 class of interneurons ^35,48^ (Figure 2A, Sup. Figure 2E, 3E). We identified 11 TN1A neurons in each hemisphere of the MANC EM volume ^40^. Our TN1A driver line targets 6.5 ± 1.38 (n=6) neurons, which stochastic labelling ^49^ confirmed are exclusively of the TN1A subtype (Sup. Figure 1F). The loss of sine song upon silencing these neurons is consistent with chronic silencing using a different driver line for TN1A neurons ^35^. The sine song elicited by activation of TN1A in isolated males persisted throughout the 10 s stimulus period (Figure 2D-E).

dMS2 is a heterogeneous group of cholinergic *fru*^+^ interneurons (Figure 2F, Sup. Figure 2F, 3F), which in the MANC EM volume fall into two distinct subtypes: a bilateral subtype represented by twelve neurons and a unilateral subtype consisting of six neurons (Sup. Fig 1G). We generated five lines that each target 10-15 dMS2 neurons (Sup. Table 1, Sup. Figure 1G). Stochastic labelling ^49^ revealed that three of the five lines target both dMS2 subtypes, whereas two are specific for the bilateral subtype (Sup. Figure 1G). Optogenetic silencing with these lines variably but consistently reduced sine song without significantly affecting pulse song (Figure 2G-H, Sup. Figure 4). The reduction in sine song reached statistical significance for four of the five lines – both lines that specifically labeled the bilateral subtype and two of the three lines that labeled both subtypes. In the activation experiments, all dMS2 lines elicited normal sine trains from isolated males, which in this case were intermittent throughout the 10s stimulus period (Figure 2I-J, Sup. Figure 4). We conclude that the bilateral dMS2 subtype is central to sine song production, but we cannot rule out that the unilateral subtype also contributes to sine song production.

Some other cell types, notably dMS9, vMS12, vPR6, and vPR13 (Sup. Table 1, Sup. Figure 1-3) also elicited varying amounts of sine song upon optogenetic activation in isolated males (Sup. Figure 5-7). However, sine song was not appreciably reduced by silencing any of these neurons, and in the case of vPR6 and vPR13, activation also elicited abnormal sounds that could not be classified as either sine or pulse song (Sup. Figure 5). These neurons also did not cluster phenotypically with the TN1A and dMS2 neurons (Figure 1C). We infer that, although these neurons can trigger varying amounts of sine song when artificially activated, they are not a core part of the circuitry that generates sine song.

### Neurons that control pulse song

Acutely silencing either the pMP2, dPR1, dMS9, or vMS12 cells significantly reduced the amount of pulse song (Figure 3), which for dMS9 and dPR1 was accompanied by an increase in the amount of sine song (Sup. Figure 6). Activation of pMP2 and dPR1 in isolated and courting males consistently induced pulse song, whereas dMS9 and vMS12 activation variably increased the amounts of both pulse and sine song (Figure 3, Sup. Figure 6).

**Figure 3.**
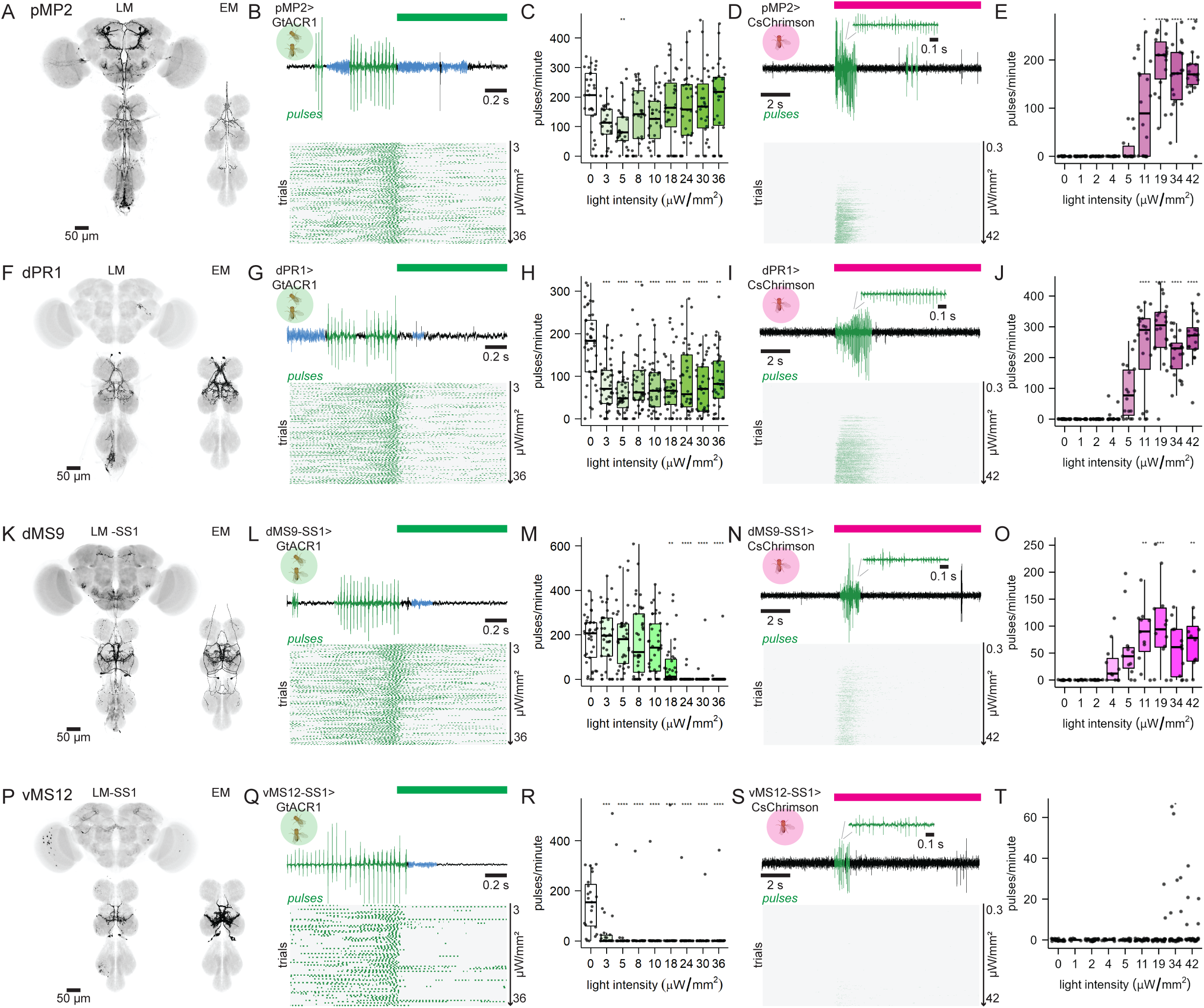
Neuron types necessary for pulse song production. A, left) A representative image of pMP2 expression. A, right) pMP2 in the EM volume. B) Response of courting pMP2>GtACR1 males to green light stimuli that were producing pulse at stimulus onset, (top) a representative example, (bottom) ethogram of pulses (green) over time with each row showing a fly/trial and rows sorted by increasing light intensity. C) Quantification of pulses/minute produced by courting pMP2>GtACR1 males without green light (0) and at increasing green light intensities. D) response of isolated pMP2>CsChrimson males to red light stimuli, (top) a representative example, (bottom) ethogram of pulses over time with each row showing a fly/trial and rows sorted by increasing light intensity. E) Quantification of pulses/minute produced by isolated pMP2>CsChrimson males without red light (0) and at increasing red light intensities. F-O) Same as A-E), but for dPR1 (F-J) and dMS9 (K-O), and vMS12 (P-T), respectively. Pulse and sine trains are indicated in green and blue, respectively. p-values calculated via Kruskal-Wallis tests and Dunn’s pairwise comparisons with Bonferroni correction. *p<0.05, **p<0.005, ***p<0.0005, ****p<0.00005.

The pMP2 neurons are a pair of cholinergic *fru*^+^ descending neurons (Figure 3A, Sup. Figure 2A, 3A). Optogenetic silencing of pMP2 tended to reduce the amount of pulse song, but this effect was significant only at one stimulus intensity (Figure 3B-C). pMP2 activation in isolated males was sufficient to drive normal pulse trains that lasted for only a few seconds after stimulus onset (Figure 3D-E). pMP2 activation in courting males increased the amount of pulse song and reduced the amount of sine song (Sup. Figure 6F-G). The pIP10 descending neurons discussed below are the primary drivers of both pulse and sine song ^31,50^, but these data suggest that the pMP2 descending neurons have a supporting role in the production of pulse song.

The dPR1 neurons are bilaterally paired cholinergic *fru*^+^ *dsx*^+^ VNC interneurons (Figure 3F, Sup. Figure 2B, 2B). Optogenetic silencing of dPR1 significantly reduced pulse song production, whereas optogenetic activation in isolated males was sufficient to produce pulse trains (Figure 3G-J). Normal pulse trains were produced for up to 5 seconds of a 10 second activation window in isolated males. Optogenetic activation during male-female courtship increased pulse song production and decreased sine song production during the activation window (Sup. Figure 6B-C). The results indicate that normal dPR1 activity is essential for pulse song and indirectly inhibits sine song production, confirming and extending previous findings from experiments using other dPR1 driver lines and chronic silencing or thermogenetic activation ^31^.

dMS9 is a previously unidentified cholinergic *fru*^+^ bilaterally paired neuron that sends ascending axon projections to the subesophageal zone in the brain (Figure 3K, Sup. Figure 2C, 3C). We derived two split-GAL4 lines that target dMS9, *dMS9-SS1* and *dMS9-SS2* (Sup. Figure 1D). Optogenetic silencing with either line significantly reduced pulse production (Figure 3L-M, Sup. Figure 6). Optogenetic activation in isolated males produced song with pulse and sine trains that lasted for up to ∼2 seconds at the beginning of a 10 second activation window (Figure 3N-O, Sup. Figure 6). These song bouts included pulse and sine song trains with largely normal characteristics, but also included abnormal pulses with three or more cycles per pulse (polycyclic pulses, Sup. Figure 6I-J). Optogenetic activation of dMS9-SS1 increased pulse song, while dMS9-SS2 increased sine song in courting males (Sup. Figure 6M, T). We conclude that dMS9 is essential for pulse song production and may play a supporting role in sine song.

vMS12 is a heterogenous group of cholinergic *fru*^+^ interneurons (Figure 3P, Sup. Figure 2D, 3D). vMS12 consists of at least two groups of anatomically distinct cell types: one group arborizes bilaterally while the other group arborizes largely ipsilaterally (Sup. Figure 1E). EM analysis identified 14 bilateral and 6 unilateral vMS12 neurons. We generated three split-GAL4 lines that each target 7-9 vMS12 neurons (Sup. Table 1, Sup. Figure 1E). Stochastic labelling ^49^ revealed that two of the lines (*vMS12-SS1* and *-SS2*) label both uni- and bilateral vMS12 subtypes, while the third line (*vMS12-SS3*) targets only the bilateral subtype. Both the *vMS12-SS1* and *-SS2* lines, but not *vMS12-SS3*, reduced the amount of pulse song in silencing experiments (Figure 3Q-R, Sup. Figure 6). The *vMS12-SS3* line also did not phenotypically cluster with the other vMS12 and dMS9 lines (Figure 1), suggesting that the unilateral vMS12 subtype may be principally involved in pulse song. None of these lines consistently increased the amount of pulse song in activation experiments, although short bouts of pulse song were elicited in isolated males at some stimulus intensities with *vMS12-SS1* (Figure 3S-T, Sup. Figure 6). Optogenetic activation of *vMS12-SS1* and *-SS2* increased sine song in courting males (Sup. Figure 6). The results indicate that vMS12 is required for pulse song production, and may also have a supporting role in sine song.

Two other cell types, dMS5 and vPR6, produced pulse-like song when activated in isolated males, but neither cell type strongly affected pulse or sine song in silencing assays (Sup. Figure 5, 7A-B). Furthermore, for vPR6, activation in courting males tended to reduce rather than increase pulse song production. Accordingly, we do not consider either cell type to be a core part of the pulse song circuit.

### Multimodal song neurons

Optogenetic silencing or activation of two cell types, pIP10 and vPR9, consistently altered the amounts of both pulse and sine song. The pIP10 neurons are bilaterally paired cholinergic *fru*^+^ neurons that receive input from neurons that integrate courtship-related auditory and visual input in the brain and make most of their excitatory synaptic outputs in the VNC wing neuropil (Figure 4A) ^30,31,50^. pIP10 activity has been shown to be necessary for pulse and sine song production and to play a state-dependent role in influencing song choice ^9,31,50,51^. In agreement with this previous work, we found that optogenetic silencing of pIP10 acutely and significantly reduced the amounts of both pulse and sine song (Figure 4B-C, F-G). Activation of pIP10 in isolated males elicited normal pulse trains throughout the stimulus duration of pIP10 activation, often interspersed with short bouts of sine song (Figure 4D-E, H-I). In courting males, pIP10 activation at low stimulus intensities increased both pulse and sine song production, but at higher stimulus intensities significantly increased pulse song production at the expense of sine song (Sup. Figure 7C-D).

**Figure 4.**
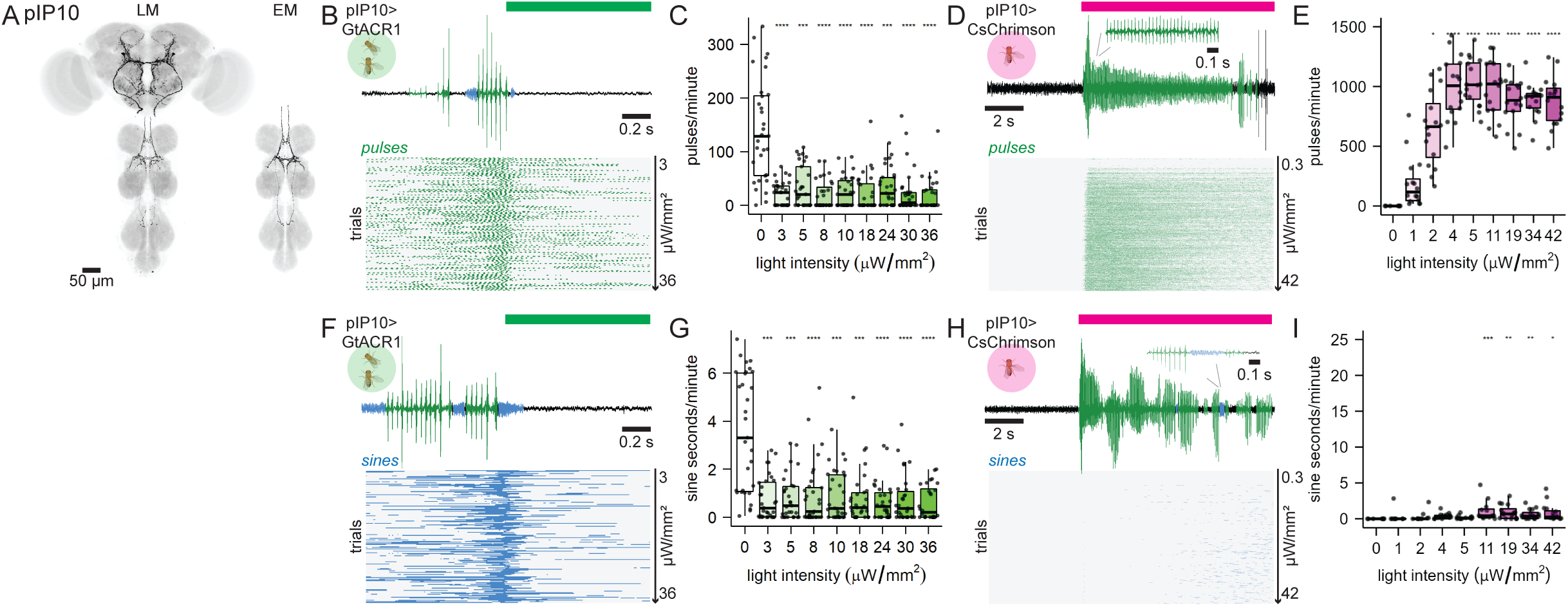
Descending control of song production via pIP10. A, left) A representative image of pIP10-GAL4 expression. A, right) pIP10 in the EM volume. B) Response of courting pIP10>GtACR1 males to green light stimuli that were producing pulse at stimulus onset, (top) a representative example, (bottom) ethogram of pulses (green) over time with each row showing a fly/trial and rows sorted by increasing light intensity. C) Quantification of pulses/minute produced by courting pIP10>GtACR1 males without green light (0) and at increasing green light intensities. D) Response of isolated pIP10>CsChrimson males to red light stimuli, (top) a representative example, (bottom) ethogram of pulses over time with each row showing a fly/trial and rows sorted by increasing light intensity. E) Quantification of pulses/minute produced by isolated pIP10>CsChrimson males without red light (0) and at increasing red light intensities. Pulse and sine trains are indicated in green and blue, respectively. p-values calculated via Kruskal-Wallis tests and Dunn’s pairwise comparisons with Bonferroni correction. *p<0.05, **p<0.005,***p<0.0005, ****p<0.00005.

vPR9 is a group of *fru*^+^ GABAergic VNC interneurons located at the midline (Figure 5, Sup. Figure 2G, 3G). We identified 7 vPR9 neurons in the EM volume and generated 5 lines that target 1-4 vPR9 neurons each (Sup. Figure 1H). Activating vPR9 in courting males using any of these lines significantly reduced one or both song types: activation using *vPR9-SS1* and *-SS2* (1-2 vPR9/animal) preferentially reduced pulse song (Figure 5B-I), *vPR9-SS3* and *vPR9-SS4* (2-3 vPR9/animal) significantly reduced sine but not pulse song (Figure 5J-Q), and *vPR9-SS5* (3-4 vPR9/animal) significantly reduced both pulse and sine song (Figure 5R-U). Silencing vPR9 using *vPR9-SS5* significantly reduced the amount of both pulse and sine song (Figure 5V-X), whereas none of the other vPR9 drivers significantly modified the amount of song upon silencing (Sup. Figure 7E-L). We conclude that normal activity among the population of vPR9 neurons is required to produce both song types and that individual vPR9 neurons may have distinct roles in pulse and sine song production.

**Figure 5.**
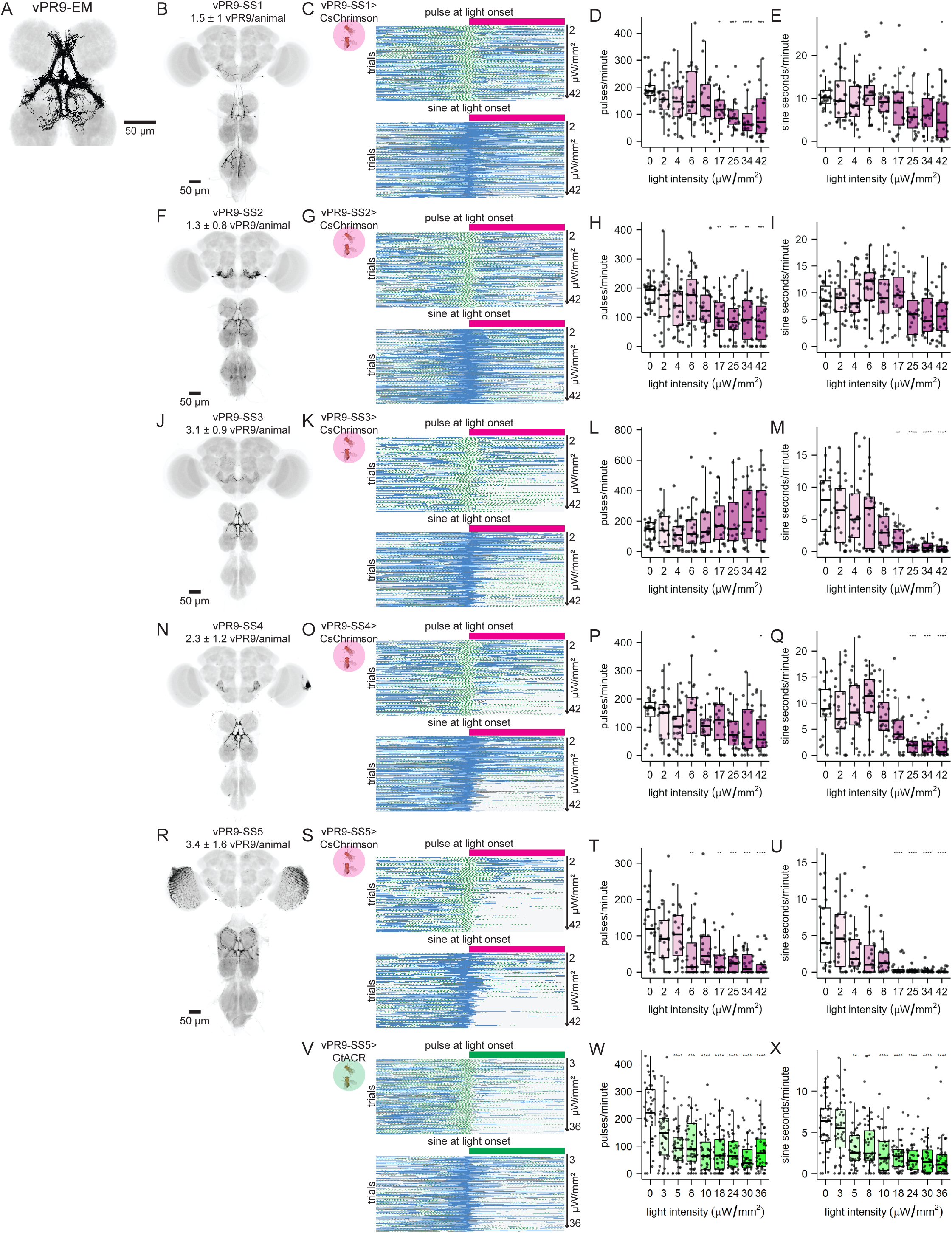
Role of vPR9 in song production. A) vPR9 neurons identified in EM. B) A representative image of vPR9-SS1-GAL4 expression. C) Response of courting vPR9-SS1>CsChrimson males to red light stimuli that were producing either pulse (top) or sine (bottom) at stimulus onset, ethograms of pulses (green) and sine (blue) over time with each row showing a fly/trial and rows sorted by increasing light intensity. D) Quantification of pulses/minute produced by courting vPR9-SS1>CsChrimson males without red light (0) and at increasing red light intensities. E) Quantification of sine seconds/minute produced by courting vPR9-SS1>CsChrimson males without red light (0) and at increasing red light intensities. F-U) Same as B-E, but for vPR9-SS2 (F-I), -SS3 (J-M), -SS4 (N-Q), and -SS5 (R-U). V-X) Same as R-U, but for GtACR1 silencing and green light. p-values calculated via Kruskal-Wallis tests and Dunn’s pairwise comparisons with Bonferroni correction. *p<0.05, **p<0.005, ***p<0.0005, ****p<0.00005.

To investigate the role of vPR9 in song production further, we imaged vPR9 activity in singing flies by monitoring GCaMP in *vPR9-SS1* and *-SS3* (Figure 6). vPR9 neurons in both lines were active during pulse and sine song production. *vPR9-SS1* neurons were more active during pulse than sine song, reflecting their preferential impact on pulse song production upon activation (Figure 6A-B). The vPR9 neurons labeled by *vPR9-SS3*, in contrast, were equally active during pulse and sine song (Figure 6C-D). This pattern has also been observed for the sine-promoting TN1A neurons, consistent with the view that sine song is produced in part by activating neurons that are also active during pulse song ^48^.

**Figure 6.**
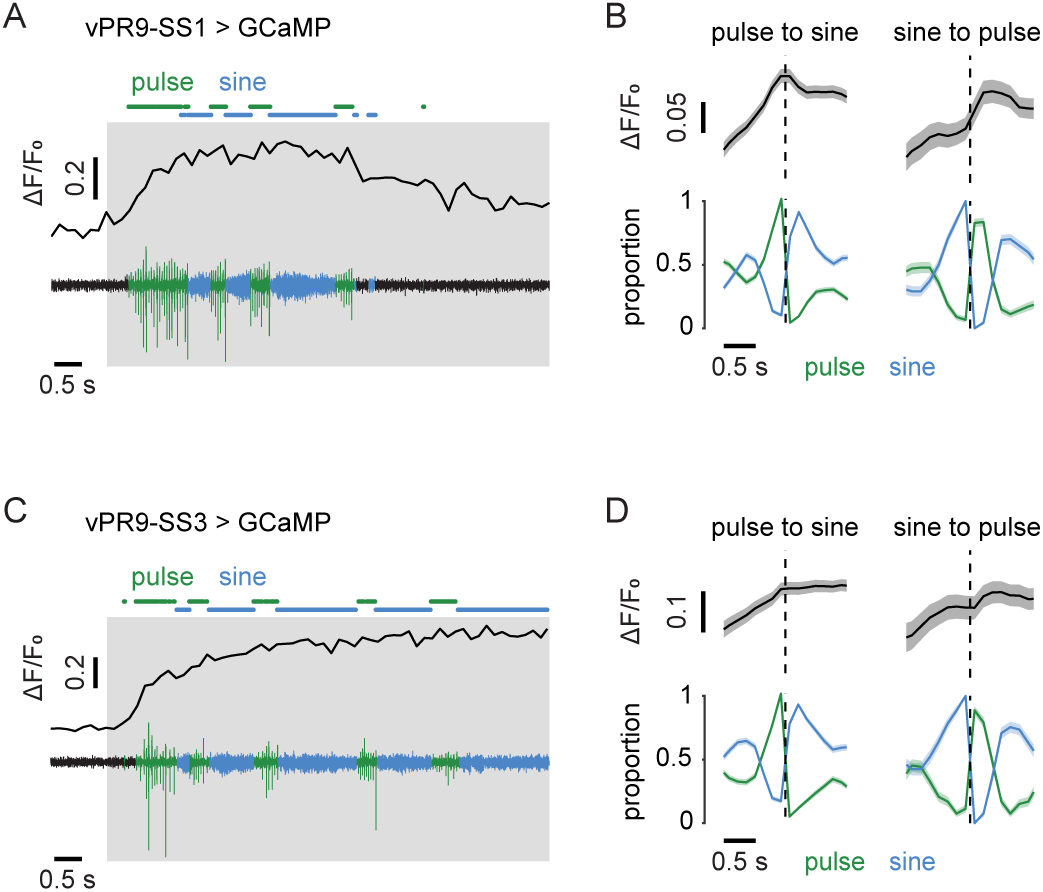
Imaging vPR9 calcium activity during song production. A) Example ΔF/F0 of neuron arbors labeled by vPR9-SS1 recorded simultaneously in a single individual (top) together with the sound recording (bottom). Song was driven by optogenetic activation of courtship promoting neurons in the brain (grey box). B) ΔF/F0 of vPR9-SS1 neurons (top) and song (bottom) around song-type transitions. Dashed vertical lines represent the transition time. Mean ± SEM across transitions for both ΔF/F0 and song. C-D) Same as (A-B), but for vPR9-SS3. vPR9-SS1 data are from 7 flies (N = 384 and 134 events for pulse-to-sine and sine-to-pulse transitions, respectively). vPR9-SS3 data are from 5 flies (N = 341 and 106 events for pulse-to-sine and sine-to-pulse transitions, respectively).

### Neurons that shape specific song features

In addition to the amounts of pulse and sine song produced in our experiments, we also quantified six song features: the number of cycles per pulse, pulse carrier frequency, inter-pulse interval, pulse amplitude, sine carrier frequency, and sine amplitude (Figures 1 and 7). This analysis indicated that the eight neuron types required to produce normal amounts of song also contribute to every feature of song and these contributions were not necessarily restricted to the primary song mode in which they function (Figure 7, Sup. Figure 8-11). For example, silencing or activating the “sine” neurons TN1A and dMS2 not only affected sine song but also some features of pulse song, such as the pulse amplitude, carrier frequency, or number of cycles per pulse (Figure 7, Sup. Figure 8). Conversely, manipulations of the “pulse” neurons dMS9 and vMS12 not only affected pulse frequency and amplitude, but also the sine amplitude or frequency (Figure 7, Sup. Figure 9-10). Silencing or activating either of the “multimodal” neurons pIP10 or vPR9 similarly affected multiple features of both song modalities (Figure 7, Sup. Figure 9, 11). These findings further strengthen the hypothesis that pulse and sine song are distinct outputs of a mostly shared circuit, rather than being generated by distinct motor circuits.

**Figure 7.**
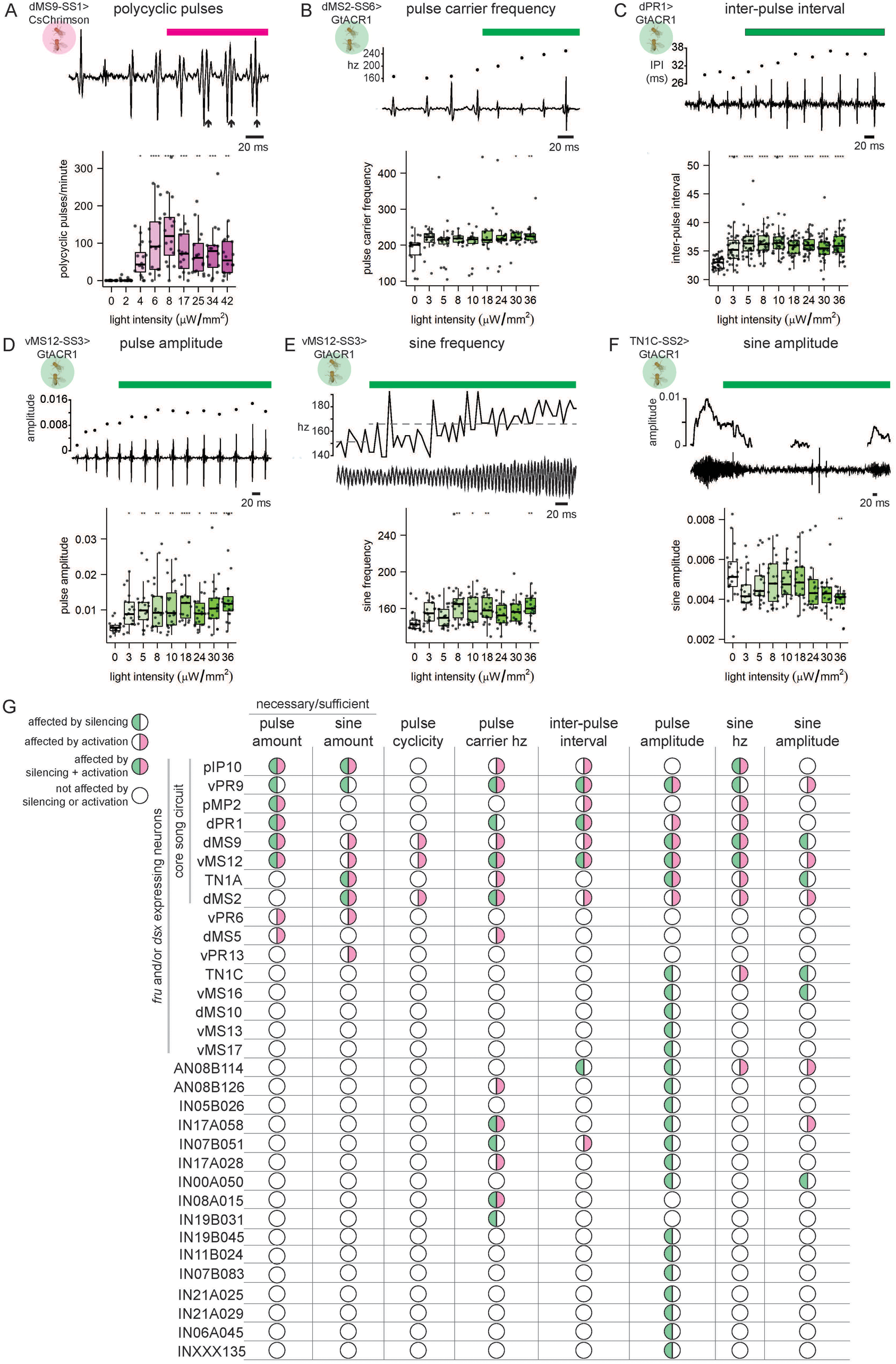
Summary song neuron activation and silencing phenotypes. A-F) Example (top) and quantification (bottom) of song features produced by courting males that were modified by optogenetic neuronal activation or silencing. One representative neuron type and activation or silencing phenotype shown for each song feature. A) Polycyclic pulses (arrows) indicated. B) Carrier frequency of each pulse is plotted. C) Each inter-pulse interval is plotted. D) Amplitude of each pulse is plotted. E) Instantaneous and average (dotted line) sine frequency plotted. F) Sine amplitude plotted. G) Summary of optogenetic silencing (green) and activation (magenta) phenotypes that affect the amount of song produced or song features. “Necessary/sufficient” phenotypes shown are silencing phenotypes in courting males that reduced the amount of song produced (green) and activation phenotypes in isolated males that induced song production (magenta), respectively. Silencing phenotypes that increased the amount of song produced in courting males (i.e., dPR1 and dMS9 silencing increased sine song) and activation phenotypes that modified the amount of song produced in courting males are not represented. Song feature phenotypes are those that occurred by silencing and/or activation of neurons in courting males. Tested neuron types with no significant effect on song are not shown. Box plots show individual animals (circles) and mean ± SD. Box plot color relates to light optogenetic stimulus wavelength and intensity. p-values calculated via Kruskal-Wallis tests and Dunn’s pairwise comparisons with Bonferroni correction. *p<0.05, **p<0.005, ***p<0.0005, ****p<0.00005.

Of particular interest was the finding that silencing or activating any of the three cell types vPR9, dPR1, or vMS12 altered the inter-pulse interval (Figure 7G, Sup. Figure 9-11). The inter-pulse interval encodes species identity, and neurons in the female brain that function in sexual receptivity are tuned to the *D*. *melanogaster* inter-pulse interval of ∼35ms ^13^. While it remains unclear exactly how the inter-pulse interval is generated, the neurons identified here may contribute to the circuit that encodes inter-pulse interval, which is critical species-specific auditory information used during courtship.

Neurons outside the core set of eight cell types rarely altered specific song features and, in most cases, altered song features only in either the silencing or the activation experiments (Figure 7, Sup. Figure 12). While many of these effects might be indirect, some may be worth investigating further. For example, the IN17A058 and IN08A015 neurons altered the pulse carrier frequency both when silenced and when activated, suggesting a direct role in song patterning. Additionally, AN08B114 is the only non-core neuron that altered the inter-pulse interval when silenced, a song parameter of particular biological importance ^12,52,53^.

### Structural organization of the song circuit

Having functionally characterized many of the individual song circuit neurons, we next examined their patterns of connectivity with each other and with wing motor neurons ^41^ within the MANC connectome^40^. This analysis revealed that the eight neuron types required to produce normal amounts of pulse and sine song (pIP10, pMP2, dPR1, dMS9, vMS12, TN1A, dMS2, and vPR9) are highly interconnected to each other and to wing motor neurons (Figures 8A-B, Sup. Fig 13). There are also connections among this group of neurons with other tested *fru*^+^ neuron types, but relatively few to the tested *fru*^-^ neuron types, or between the non-core *fru*^+^ and *fru*^-^ types (Figure 8A). Overall, the number of connections to wing motor neurons is comparable for both the *fru*^+^ and *fru*^-^ sets. This broad pattern of connectivity supports the view that the song-generating circuitry comprises the eight core *fru*^+^ cell types, likely augmented by several other *fru*^+^ cell types, and that the song circuit is largely distinct from the motor circuits for flight and for other behaviors that also require wing motor neuron activity.

**Figure 8.**
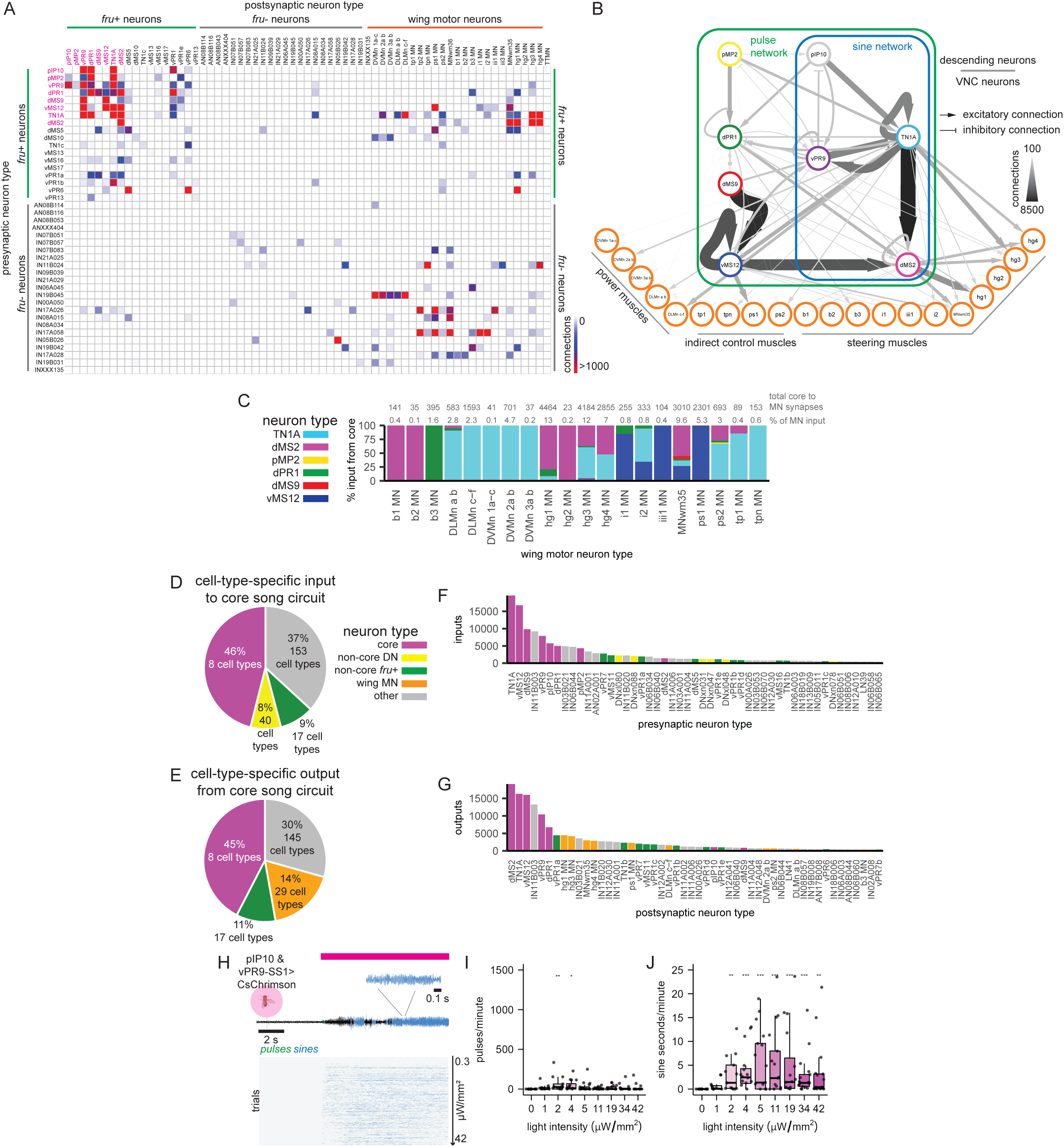
Song circuit structure. A) Connections among functionally tested neurons and wing motor neurons by neuron type. Core song circuit neurons are highlighted in magenta. B) Wiring diagram of the core song circuit and wing motor neurons. Pulse and sine networks are highlighted. C) Core circuit connectivity to wing motor neurons. Percentage of total input to each motor neuron from the core circuit is plotted. Total connections from the core circuit to each motor neuron are shown in grey above. D) Cell-type specific input to the core song circuit neurons as a percentage of the total input to the core song neurons. Cell types are grouped into one of four indicated categories. E) Same as (D), but output from core song neurons. F) Top 50 inputs to the core song circuit ranked by number of connections. G) Same as (F), but for output from the core song circuit. C-G) Threshold of 10 connections/neuron pair. H-J) Response of isolated pIP10+vPR9-SS1>CsChrimson males to red light stimuli. I) (top) a representative example, (bottom) ethogram of pulses (green) and sine (blue) over time with each row showing a fly/trial and rows sorted by increasing light intensity. I-J) Quantification of (I) pulses/minute and sine (J) seconds/minute produced by isolated pIP10+vPR9-SS1>CsChrimson males without red light (0) and at increasing red light intensities. p-values calculated via Kruskal-Wallis tests and Dunn’s pairwise comparisons with Bonferroni correction. *p<0.05,**p<0.005, ***p<0.0005, ****p<0.00005.

Within the core song circuit, there are two largely feed-forward excitatory networks from descending input to the motor output that drive each song type (Figure 8B, Sup. Figure 14A). The feed-forward sine network consists of the three neuron types that are necessary and sufficient to produce sine song: pIP10, TN1A, and dMS2. The feed-forward pulse network includes all members of the feed-forward sine network (pIP10, TN1A, and dMS2) plus pMP2, dPR1, dMS9, and vMS12. There are also strong recurrent connections in the core song circuit that are embedded within these feed-forward networks (Figure 8B, Sup. Figure 13, 14B). TN1A makes strong recurrent connections with dPR1. Additionally, the inhibitory vPR9 neuron is not part of the feed-forward networks but is a member of the core song circuit and makes strong recurrent connections with dPR1, TN1A, and pIP10. Further, TN1A, vPR9, vMS12, and dMS2 make strong recurrent connections within their respective populations of neurons. dPR1, TN1A, and vMS12 also make strong connections between the left and right neurons within each cell type (Sup. Figure 14C).

### Song circuit connections with wing motor neurons

*Drosophila* wing movements are controlled by three sets of muscles. The power muscles consist of the dorsal longitudinal muscles (DLMs) and dorsal ventral muscles (DVMs), which are positioned orthogonally to each other and divided into six and seven large fibers, respectively ^54^. One motor neuron innervates each of these fibers, but the neurons do not fire one-for-one with muscle contraction. Instead, they fire asynchronously to induce contraction, which deforms the thorax, thereby stretch-activating the opposing muscles and producing the fast wing beat frequencies observed during flight and courtship song ^54^. The second set of muscles consist of 12 direct control muscles (or steering muscles; b1-b3, i1, i2, iii1, iii3, iii4, hg1-hg4), which regulate wing extension and fine wing movements ^39,54^. Each of these muscles is innervated by a single motor neuron that fires one-for-one with muscle contraction. The third set of muscles includes five indirect control muscles (tp1, tp2, ps1, ps2, tt) that regulate thorax resonance ^54^. One motor neuron innervates each of these muscles, with the exception of tp1 and tp2 muscles which receive additional input from the tpn motor neuron ^39,55^. Motor neurons fire one-for-one with indirect control muscle contraction.

Electrophysiological recordings from power muscles have demonstrated that these muscle fibers are sporadically but rhythmically active during both pulse and sine song production ^36,38^. Most direct core song circuit output to the power muscle (DLMs and DVMs) motor neurons is made by TN1A neurons, but there are also connections from the “pulse” neurons dPR1 (Figure 8D, Sup. Figure 13). Both pulse and sine feed-forward networks include strong connections to TN1A and silencing a subset of TN1A neurons significantly inhibited the amount of sine song produced and modulated pulse and sine song features (Figure 4, 7). Calcium imaging experiments further demonstrate that TN1A neurons are active during both pulse and sine song production ^48^. We conclude that TN1A serves as the primary output of the core song circuit to the power muscles to drive fast wing oscillations during sine song, with additional outputs such as dPR1 being recruited during pulse song.

Calcium imaging of the wing muscles during song production has demonstrated that the direct control muscles i1, hg1-4, and b3 are all strongly active during pulse song production ^39^. iii3, iii4, and i2 were either active during pulse song or not on a trial-by-trial basis, and iii1 demonstrated weak activity during pulse song bouts. Further, i1 and iii1 were preferentially active during pulse song production compared to sine song. Similarly, b3, hg2, hg3, and hg4 may be weakly preferentially active during pulse song ^39^. Connectivity to these direct control motor neurons from the core song circuit neurons is largely consistent with these observations (Figure 8D, Sup. Figure 13). All the associated motor neurons except iii3 receive direct input from the core song circuit. Additionally, i1, iii1, and b3 MNs are exclusively innervated by core neurons that were necessary to produce pulse but not sine song (Figure 8D, Sup. Figure 13). Conversely, hg1-hg4, i2, and MNwm35 (likely iii4 ^41^) are innervated by a combination of core song neurons with the strongest connections coming from dMS2 and TN1A. Finally, consistent with the observation that b1 and b2 muscles are silent during song ^39^, there are no strong connections from the core song circuit to the b1 or b2 motor neurons (Figure 8D).

There are few clear recordings of the indirect muscles during song production. Electrical recordings from possible tp1 and tp2 muscles suggest that they are active during the wing stroke of pulse and sine song (^37^; called b3 and b4). tp2 motor neuron silencing phenotypes are currently inconsistent across studies ^39,56^, but silencing the tpn motor neurons was shown to abolish pulse and sine song ^39^. There are no direct connections from the core song circuit to tp2, but tp1 and tpn are weakly innervated by TN1A and dMS2. Silencing ps1 motor neurons produces pulse-specific phenotypes ^34,39,56^. Consistent with this, the ps1 motor neurons are innervated almost exclusively by vMS12 (Figure 8D, Sup. Figure 13), which is necessary to produce pulse but not sine song. The modified pulse frequency we observed upon acute silencing of vMS12 (Sup. Figure 10) also resembles the observed effect of chronic silencing of ps1 motor neurons ^34,39^.

In total, the connectivity between the core song circuit and the wing motor neurons is consistent with published accounts of wing muscle and motor neuron activity during song production. It is also consistent with the view that neurons necessary for sine song are also active and function during pulse song. Notably, the “sine” neurons TN1A and dMS2 make most of the direct connections from the core song circuit to wing motor neurons critical for both pulse and sine song production (e.g., the power muscles, hg1-4), whereas the “pulse” neurons dPR1 and vMS12 provide the major input from the song circuit to the motor neurons and muscles used preferentially during pulse song (ps1, i1, iii1, b3).

### A possible circuit mechanism for switching between pulse and sine song

Collectively, the eight core song cell types we have identified receive almost half of their synaptic input from other members of this group (Figure 8E), and similarly provide almost half of their synaptic output to other core neurons (Figure 8F). The strong interconnectivity amongst this group of neurons suggests that other neurons with similarly strong connections to members of this group might also function in generating or patterning courtship song. Accordingly, we ranked all the neurons with synaptic connections to this group by their total number of inputs to or outputs from core song circuit neurons (Figure 8G-H, Sup. Figure 13).

Three uncharacterized neurons stood out in this analysis, all of which are predicted to be inhibitory (Sup. Figure 14D). Two of them, IN11B003 and IN03B021, make strong reciprocal connections with core song neurons, similar to those of the inhibitory vPR9 neurons; the third, IN06B044, is located within the feed-forward network for pulse song (Sup. Figure 14F-H). These inhibitory neurons may contribute to the generation of rhythmic oscillations, prevent runaway excitation in the feed-forward circuits, or mediate switching between pulse and sine song.

Additionally, among the inputs to the core song circuit are eight uncharacterized descending neuron types (Sup. Figure 14E). DNxl080 and DNxl048 make relatively strong connections to both the sine-promoting TN1A and pulse-promoting dPR1 neurons, while DNxl058 and DNfl038 make strong excitatory connections to dPR1 but not to TN1A (Sup. Figure 13, 14F-G). In contrast, DNxn031, DNxn088, DNxn078, and the *fru^+^* pIP9 make strong excitatory connections to the TN1A and vPR9 neurons but not to dPR1 (Sup. Figure 13, 14F-G). The vPR9 neurons, in turn, provide inhibitory input to the dPR1 “pulse” neurons, a subset of the TN1A “sine” neurons, and presynaptic inhibition to the pIP10 descending “command” neurons for song. This collection of uncharacterized DNs thus provide direct excitation and indirect inhibition of the pulse and sine feed-forward networks, suggesting that they might mediate a switch between song types.

Testing this hypothesis will require genetic access to these descending neurons. We do however have genetic access to one of the top targets of the descending neurons, vPR9, enabling us test one prediction of this model: namely, that, whereas pIP10 activation alone elicits more pulse song than sine song (Figure 4), coactivation of vPR9 should increase the ratio of sine song to pulse song upon pIP10 activation. We tested this prediction using the *vPR9-SS1* driver and found that coactivation of pIP10 and vPR9 indeed results in a courtship song composed almost entirely of sine song (Figure 8I-K). This result supports the idea that alternative descending inputs from the brain mediate switching of song modality by preferentially activating and inhibiting either the pulse or the sine feed-forward networks, likely in response to sensory cues that signal the female’s position and response ^8^.

## Discussion

In this study we have combined cell-type specific optogenetics with connectome analysis to identify key components of the neural circuits in the male VNC that generate courtship song. Our functional and anatomical analysis defines a core song circuit composed of at least eight highly interconnected cell types. All of these neurons express either *fru* or *dsx* and are therefore likely to be sexually dimorphic. They receive descending input from brain neurons known to control courtship song and provide output to motor neurons known to be active during singing. Our functional analysis identified several other cell types that also contribute to song patterning, and our anatomical analysis identified a handful of functionally untested cell types that may be an integral part of the core song circuit. Having identified and functionally characterized at least part of the core song circuit, we can begin to speculate how it is organized, how it generates a richly structured song, and how it may have evolved.

### Pulse and sine circuits

The core song circuit appears to be organized into two feed-forward pathways from descending neurons to wing motor neurons. The smaller of these networks consists of pIP10, TN1A, dMS2, and vPR9 and produces sine song. This sine song circuit is embedded within a larger network that additionally includes the remaining core song neurons, pMP2, dPR1, dMS9, and vMS12, and produces pulse song. This organization suggests that the song circuit functions in two main states: a sine state, in which only the sine neurons are active, and a pulse state, in which the additional core neurons also become active. This model, derived from optogenetic manipulations and connectome analysis, is further supported by calcium imaging experiments demonstrating that pIP10, vPR9, and TN1A are active during both pulse and sine song whereas pMP2 and dPR1 are active only during pulse song (this study and ^48^).

This proposed circuit organization shares commonalities with multifunctional rhythmic motor circuits that have been studied in other species, such as the circuits that control decapod crustacean feeding ^57^, mammalian respiration ^58,59^, and leech locomotion ^60^. The distinct rhythmic motor patterns involved in decapod feeding, mammalian respiration, and leech locomotion are generated by circuits, which, like the *Drosophila* song circuit, contain both uni- and multifunctional neurons. Furthermore, voltage imaging studies in the leech have demonstrated that more than two times as many cells are active during crawling than during swimming and that most of the cells active during swimming were also active during crawling ^60^. Similarly, for *Drosophila* song, we propose that more cells are active during pulse than during sine song, and that all of the VNC sine neurons are also active during pulse song.

### Composing the song

During song, the male switches rapidly between bouts of pulse song and bouts of sine song, each typically lasting from tens to hundreds of milliseconds. The choice between these two modes, the choice of wing used for singing, and the amplitude of song are associated with female behavior ^8,10^. The position, distance, velocity, and behavior of the female are all presumably calculated in the brain and relayed to the VNC song circuits via descending neurons. The best-studied descending input to the song circuit is the *fru*^+^ pIP10 neuron ^9,31,50,51^. pIP10 activity is necessary to produce both pulse and sine song but activating this neuron type in isolation drives pulse song with relatively little sine song. Thus, other descending neurons must also contribute to song production, in particular to sine song. We have identified in the MANC connectome several other classes of descending neurons - pIP9, DNxn031, DNxn078, and Dnxn088 - which make relatively strong input to the sine feed-forward network and little or no direct input to the pulse feed-forward network. Genetic reagents are not yet available to functionally characterize these descending neurons, but a strong prediction from their connectivity is that one or more of these neurons drives sine song. Additionally, we have functionally characterized the *fru*^+^ pMP2 neurons, which preferentially target the pulse network and whose activity is necessary and sufficient for pulse but not sine song. We propose therefore that the choice to sing and the choice of song mode are both made in the brain and communicated to the song motor circuits in the VNC via the differential activation of a set of descending neurons. A similar organization has been proposed for the choice to move and the choice between swimming and crawling in the leech ^61^.

How might this switch be implemented in the VNC? In part, the activation of either sine or pulse circuits is likely attributed feed-forward excitation from the relevant descending neurons. We note, for example, that pIP10 and pMP2 are more strongly connected to the pulse network than the sine network, whereas the converse is true for pIP9 and the other putative sine-promoting descending neurons. It seems unlikely however that the switch between pulse and sine song is mediated entirely by feed-forward excitation from descending neurons. When the switch occurs naturally, it is abrupt and complete. Moreover, we have found that a switch from pulse to sine can be elicited by artificially exciting the vPR9 neurons in the VNC. Inhibitory pathways are therefore likely to exist within the VNC to silence the pulse circuit during production of sine song and possibly to alter the activity of the “sine” neurons during pulse song. vPR9 neurons are likely a key part of these inhibitory circuits. They receive direct input from the putative sine-promoting descending neurons and inhibit the pulse-promoting dPR1 neurons. Furthermore, their activity is sufficient to switch pIP10-induced song from pulse to sine. vPR9 activity would therefore appear to be a mediator of the pulse-to-sine transition. However, confusingly, the same vPR9 neurons also receive input from pIP10 and from the pulse-promoting pMP2 and dPR1 neurons – an observation that is not readily integrated into such a simple switch model. We also note that the vPR9 neurons are an anatomically heterogeneous cell population, and that this heterogeneity may be reflected in the divergent silencing and activation phenotypes we observed using different vPR9 drivers. Genetic reagents that label specific subtypes of vPR9 and other inhibitory cell types, together with functional imaging during natural mode switches, will be required to further explore how the VNC motor circuits switch between pulse and sine song.

### Song Evolution

Courtship song has evolved extensively across *Drosophila* species. Most species produce at least one form of pulse song, whereas sine song production is far less common, and phylogenetic evidence suggests that pulse song is the ancestral song type in the genus ^62^. Furthermore, the distribution of wing muscle sexual dimorphisms across the *D. melanogaster* species subgroup suggests that pulse song is ancestral to this group and that sine song evolved later ^63^. This suggests that evolution adapted a subset of the ancestral pulse song circuit to create an additional song modality, sine song. The evolution of abundant sine song production in some species may be due to changes in connectivity between the pulse and sine networks, changes in the sine feed-forward network, or changes in the descending input to these networks. Here, too, there are potential parallels to leech locomotion ^64^. In both fly song and leech locomotion, the larger network appears to generate the ancestral mode (pulse or crawl), with only part of this network active during production of an evolutionarily derived motor pattern (sine or swim).

The specific features of pulse and sine song are highly variable across species; every *Drosophila* species that sings produces a unique song with species-specific song features ^62^. At least some of these features are critically important for species recognition and may modulate the female decision to mate with a male ^12,52,53^. We have identified candidate nodes within the song circuit that may have evolved to generate species differences in features such as the inter-pulse interval, sine and pulse frequency, and the number of cycles per pulse. Testing hypotheses about the circuit changes that underlie song evolution will require comparison of the function of homologous neurons across species. Split-GAL4 enhancer combinations identified in *D. melanogaster* have been used to target pIP10 in *D. yakuba* ^50^, suggesting that the split-GAL4 reagents we have identified may be useful for studying homologous circuits across *Drosophila* species. Testing anatomical hypotheses will require comparative connectomics, which should be facilitated by recent advances in EM ^40,65–72^ and in expansion microscopy ^30^.

### Conclusions and future directions

Our systematic functional analysis of a large number of neural cell types using quantitative optogenetic behavioral analyses combined with connectome analysis provides a detailed map of the song circuit in the male ventral nerve cord. This work provides strong hypotheses about how the song circuit contributes to the complex motor output of courtship song, and suggests common themes in the organization and evolution of multifunctional motor systems across a wide range of animal species.

Nonetheless, our study has important limitations. Connectomes reconstructed through electron microscopy lack molecular detail. For example, they do not currently reveal electrical connections or neuropeptides, both of which are major contributors to neuronal communication and circuit function. Neurons with a comparatively low synaptic count to volume ratio are suspected of communicating largely through electrical connections, and amongst the more prominent of these neurons in the male VNC are the dMS2 song neurons ^41^. At present, connectomic analysis is also typically limited to very small sample sizes, usually just one, and so we cannot address variability both within and between species. These limitations will eventually be overcome through advances in electron ^40,41,67,68,71–73^ and light ^30^ microscopy to provide molecular detail and to allow study of much larger sample sizes.

Optogenetics, like genetics more generally, provides insight into the contributions of individual components (neurons or genes, respectively) by breaking these components. Complementary information about neural circuit function can be provided by direct observation of neural activity while animals perform each motor action. Fortunately, it is now possible to drive naturalistic song in male flies while imaging circuit activity in both the brain and VNC in vivo ^32,48^. Electrophysiology, which has been critical in studies of the rhythmic motor circuits in crustaceans, sea slugs and the leech, will be considerably more challenging for the VNC *Drosophila* song circuit due to the very wing vibrations it seeks to explain. Advances in light-based sensors that report voltage, neurotransmitters, and second messengers, together with holographic optogenetics, will likely overcome these limitations and enable the simultaneous measurement of electrical and biochemical processes across large populations of neurons.

Ultimately, a useful goal will be to construct computational models of the song circuit that incorporate anatomical detail at the cellular and molecular levels and functional data at the physiological and behavioral levels. The present work provides a foundation for these future experimental and computational studies by beginning to reveal how motor circuits within the male ventral nerve cord produce and shape a motor program critical to reproductive success.

## Acknowledgments

We thank Emily Behrman, Steve Sawtelle, Ben Arthur, and Elizabeth Kim for help with courtship song recording and analysis; Kari Klose, Christina Christoforou, Yisheng He, Mark Eddison, and Gudrun Ihrke of the Janelia Project Technical Resources Team for dissecting, labeling, and acquiring the *fru* co-labeling and EASI-FISH images; Geoffrey Meissner and the FlyLight Project Team for multi-color flipout experiments and all other light microscopy images; Hideo Otsuna for MANC neuron meshes and help with LM-EM matching; Konrad Rokicki for pre-release access to NeuronBridge; Gwyneth Card, Greg Jefferis, Stuart Berg, and the FlyEM Project Team for pre-release access to the MANC connectome; Han Cheong for MANC wing motor neuron identification; Yun Ding for valuable discussions and comments on the manuscript; Troy Shirangi for valuable discussions. This article is subject to HHMI’s Open Access to Publications policy. HHMI lab heads have previously granted a nonexclusive CC BY 4.0 license to the public and a sublicensable license to HHMI in their research articles. Pursuant to those licenses, the author-accepted manuscript of this article can be made freely available under a CC BY 4.0 license immediately upon publication.

## Author contributions

J.L.L., K.W., D.L.S., and B.J.D. conceived the study. K.W., H.M.S., and M.X. generated the split-GAL4 lines. J.L.L. conducted and analyzed behavior experiments and analyzed all LM data. H.M.S. conducted and analyzed calcium imaging experiments. J.L.L. and K.W. conducted LM-EM matching and EM circuit analysis. J.L.L. produced all figures with input from coauthors. J.L.L. and B.JD. wrote the manuscript with input from coauthors. B.J.D. and D.L.S. acquired funding.

## Methods

### Experimental animals

*D. melanogaster* flies used in non-optogenetic experiments were crossed and raised on standard cornmeal-agar-based medium at 25°C with relative humidity of ∼50% and a 12:12 light/dark cycle. Flies used in optogenetic behavior experiments were crossed and raised in the dark on media that was supplemented with 0.4 mM trans-retinal. Detailed information on fly sex, housing, and age for each experiment is indicated in the relevant section below. Detailed fly genotype information is in Sup. Table 1.

### Genetic reagents

We generated sparse, split-GAL4, transgenic fly lines to target song neuron candidates. Split-GAL4 lines used in this study have p65ADZp and ZpGAL4DBD inserted at the *attP40* site and *attP2* site, respectively ^74,75^, except for lines that use *dsx*-ZpGAL4DBD as the DBD portion of the split half (Sup. Table 1) which have ZpGAL4DBD inserted at the first coding exon of *dsx* ^35^. p65ADZp and ZpGAL4DBD lines labelling neurons of interest were identified using manual inspection or using the automated color depth MIP (maximum intensity projection) mask search method ^76^. The expression of selected combinations of p65ADZp and ZpGAL4DBD was then examined with a *UAS* reporter (*20×UAS-CsChrimson-mVenus* in *attP18*) by immunofluorescence staining and confocal microscopy (https://www.janelia.org/project-team/flylight/protocols). Finally, the combinations of p65ADZp and ZpGAL4DBD that gave the most specific expression patterns were stabilized by putting the two hemi-drivers in the same flies, and *SS* (denoting sSup. Tablesplit-GAL4) numbers were assigned.

### Immunofluorescence staining

Nervous system fixation, immunohistochemistry, dehydration, and DPX mounting steps were performed using standard protocols described previously ^77^. Detailed protocols for all steps, including standard split-GAL4 screening, double-label staining, and stochastic labelling in multiple colors (MCFO) ^49^, are available at https://www.janelia.org/project-team/flylight/protocols. Briefly, brains and VNCs were dissected in Schneider’s insect medium and fixed in 2% paraformaldehyde at room temperature for 55 min. Tissues were washed in PBT (0.5% Triton X-100 in 1X phosphate buffered saline), and blocked using 5% normal goat serum before incubation with antibodies for 2-3 overnights. Tissues were then washed in PBT for 10 min 5 times before incubation in secondary antibodies for 2-3 overnights. To test if neurons targeted in split-GAL4 lines were *fru^+^*, split-GAL4 lines were crossed to a LexA line that labels *fru+* neurons ^78^ (LexAop2-Syn21-opGCaMP6s [*suHwattP8*], 10XUAS-Syn21-Chrimson88-tdT-3.1 [*attp18*]; ; fru^P^^1.LexA^), and staining was examined to determine whether the split-GAL4 and fru^P^^1.LexA^ were co-expressed in the same neurons.

### EASI-FISH

The primary neurotransmitter used by song neurons was determined using in situ hybridization. Expansion-assisted iterative fluorescence in situ hybridization (EASI-FISH) ^79,80^ was utilized and published oligonucleotide probes for acetylcholine (ChAT), GABA (Gad1), glutamate (vGlut), tyramine (Tdc2, Tβh), octopamine (Tdc2, Tβh), serotonin (SerT), and dopamine (pale) were used ^81,82^. The protocol used for EASI-FISH was identical to that described by Eddison and Ihrke, 2022.

### Light microscopy imaging and analysis

Confocal imaging of unexpanded samples was performed on a ZEISS LSM 800 or an LSM 880 inverted confocal microscope with a Plan-Apochromat 20×/0.8 M27 objective or a Plan-Apochromat 63×/1.4 oil-immersion objective (ZEISS). Images were captured using ZEN software (ZEISS). Expanded samples were imaged on a ZEISS Lightsheet 7 with a W Plan-Apochromat 20X/1.0 water-immersion objective (ZEISS). Split-GAL4 and MCFO images (e.g., Sup. Figure 1) were aligned to the JRC 2018 central brain and VNC unisex templates ^83^. Images were evaluated (Sup. Table 1) using VVDViewer and all neuron images including images of EM neuron meshes aligned to the JRC 2018 VNC unisex template (e.g., Sup. Figure 1) were captured in VVDViewer ^84^ (https://github.com/JaneliaSciComp/VVDViewer).

### Behavior experiments and analysis

For all optogenetic behavior experiments (CsChrimson and GtACR1), male flies were collected on the day of eclosion and group housed in the dark at 25 °C on standard medium supplemented with 0.4 mM trans-retinal. For paired male-female courtship experiments, male flies were paired with 1-2 day old group housed virgin females. All flies were transferred from food to chambers without anesthesia using a mouth aspirator.

Behavior experiments were conducted in 5 mm chambers in a song recording rig descried in Sawtelle et al., 2023 (in preparation). In brief, the chambers were enclosed on top by a transparent plastic cover and on the bottom by a fine mesh. Cameras placed above the chambers recorded video and a microphone was positioned under each chamber to record audio. IR light was used for video recording. Both audio and video were recorded at 5 kHz. Dim 465 nm light illuminated the chambers to allow the flies to see.

The light stimuli protocols used for all optogenetic experiments are illustrated in Figure 1B. For single male optogenetic neuronal activation experiments, individual male flies were placed into a behavioral chamber. Ten second, constant on 625 nm red light stimuli of varying light intensities were applied and interspersed by ten second intervals without a stimulus. Stimulus light intensities ranged from 0.3 to 42 µW/mm^2^, and each intensity was repeated three times. Stimuli were applied in a pseudo-random order (to ensure that the same intensity was not applied in consecutive bouts). A representative subset of the stimuli used is shown in all scatter and box plot figures. For optogenetic neuronal activation experiments in courting males, individual male flies were placed into a behavioral chamber with an individual female fly. Three second, constant on 625 nm red light stimuli of varying light intensities were applied and interspersed by ten second intervals without a stimulus. Stimulus light intensities ranged from 2 to 42 µW/mm^2^, and each intensity was repeated ten times. Stimuli were applied in a pseudo-random order. This protocol was identical for optogenetic silencing experiments in courting males, except 525 nm green light stimuli that ranged from 3 to 36 µW/mm^2^ were used in place of red light stimuli.

Courtship song was classified in audio files using SongExplorer ^47^. A convolutional neural network model was trained to detect pulses, sine waves, polycyclic pulses, inter-pulse intervals, ambient sound, and other sounds (composed largely of high amplitude wing sounds generated by flight attempts). We found that training and classifying inter-pulse intervals, ambient, and other sounds increased the accuracy of pulse, sine, and polycyclic pulse classification. Ground truth data for the model consisted of the following manually annotated song events: 1695 pulses, 424 sine waves, 1996 polycyclic pulses, 2562 inter-pulse intervals, 88 ambient waves, and 53 other sounds. The model was trained for 7 million steps until the loss, accuracy, and error rates plateaued, resulting in the following precision/recall statistics: pulse 89%/81%, sine 93%/98%, and polycyclic pulse 83%/88%. SongExplorer was used to generate ethograms of each audio recording, which was used for further song analysis.

Custom Matlab scripts ^3^ were used to extract basic song statistics from the ethograms generated via SongExplorer. These include pulses/minute, pulse trains/minute, median pulse train length, mode inter-pulse interval, mode pulse frequency, median pulse amplitude, polycyclic pulses/minute, sine seconds/minute, sine trains/minute, median sine train length, mode sine frequency, and median sine amplitude. All other song behavior statistics and plots were computed and generated via custom R scripts. To compare the effect of light stimuli on song, within experiment trials for each stimulus intensity were extracted and compiled, song statistics for each stimulus intensity were computed, and the song statistics from each stimulus intensity were compared to song statistics from periods of no optogenetic stimulus across flies using Kruskal-Wallis tests and Dunn’s pairwise comparisons with Bonferroni correction (e.g., Figure 2C). To visualize song ethograms before and after light stimuli (e.g., Figure 2B), audio clips 1 second before and after light stimuli were extracted and separated into instances when pulse, sine, or no song were being produced at light onset. To cluster fly lines in Figure 1C, we calculated the Dunn’s test z-statistic across all flies and trials for each experimental condition, and then for each line selected the maximum absolute z-statistic across all stimulus intensities. A vector of these z-statistics was generated for each fly line. The Pearson correlation coefficient among the fly line was calculated based on these vectors and the lines were clustered based on this correlation and average linkage.

### Calcium imaging

All calcium imaging and analysis methods are exactly as described in Shiozaki et al., 2022. In brief, jGCaMP7f ^85^ was imaged in the arbors of vPR9-SS1 or vPR9-SS3 neurons before, during, and after optogenetic activation of 22D03-LexAp65 via CsChrimson ^46^. Activating the brain neurons driven by 22D03-LexAp65 – which includes a subset of the courtship gating neurons, P1 – is sufficient to drive naturalistic pulse and sine song ^48^. GCaMP activity was compared when each fly was singing pulse, sine, or no song.

### EM-LM neuron matching and neuron naming

Neurons expressed in split-GAL4 lines were matched to MANC bodyIDs via manual inspection of the MANC neuron collection ^40^ and NeuronBridge ^86^. We only attempted to match the targeted wing neuropil-innervating neurons present in each split-GAL4 line and did not match any off-target expression (Sup. Table 1). Therefore, unless noted otherwise, we were attempting to match a single cell-type expressed in each split-GAL4 to anatomically similar MANC bodyIDs. In many instances our cell type definitions included multiple MANC v1.0 systematic cell types. We chose to err on the side of including bodyIDs that may not be represented by a given split-GAL4 line but could not be clearly eliminated based on light-level anatomy rather than attempting to limit each LM-defined cell type or split-GAL4 line to a single EM-defined systematic cell type.

This strategy to reduce false negative LM-EM matches is consistent with our EM analysis goals. We aimed to analyze connectivity among the tested neurons and wing motor neurons (Figure 8A-D) and identify additional neurons that are strongly connected to the core song circuit and may contribute to song production (Figure 8E-H, Sup. Figure 13-14). In the first instance, we are confident in our broad core song circuit neuron EM-LM matches (even if a given split-GAL4 may only represent a portion of the total population) and found that non-core tested neurons were weakly connected to the core – even if we include multiple systematic types for a given cell type. Therefore, it is unlikely that our decision to err on the side of false positive LM-EM matches resulted in any significant false positive connections among tested neurons and wing motor neurons. In the second instance, including additional bodyIDs in each systematic type increased our ability to identify potential core-like neurons based on the strength of connectivity with the core.

Notably, the systematic type naming scheme has been refined in subsequent versions of the MANC, which has resulted in fewer bodyIDs in many systematic types. Future work will be required to determine if the split-GAL4 lines generated here or subsequent lines targeting song neurons can be further refined to fit the cell-type definitions generated by EM analysis. Furthermore, additional work is required to identify the optimal approach for defining a neuron type.

For *fru^+^* neuron types, we used previously assigned names where applicable ^31,35,44^ and assigned names for newly identified *fru^+^* neuron types based on the previously established *fru^+^* neuron naming scheme ^44^. We named *fru^-^* neuron types for which we have split-GAL4 lines based on the matched MANC v1.0 systematic type. In cases where we identified multiple possible systematic type matches for a given split-GAL4, we used the systematic type with the best match or lowest number (e.g., AN08B116 is used for a neuron type that may include systematic types AN08B116, AN08B119, AN08B125, and AN08B126). In cases where no systematic type was assigned to the bodyID in v1.0 of the MANC, the systematic type from v1.1 was used and is indicated in Sup. Table 1. Finally, in cases where cell types described here resemble those targeted in a recently published collection of split-GAL4 reagents targeting wing neurons ^56^, we note the possible matches in Sup. Table 1.

### EM analysis

All EM analysis was performed on the MANC v1.0 volume and annotations ^40,41^. We wrote Python scripts to identify the connections between every tested neuron type bodyID, wing motor neuron bodyID ^41^, and every bodyID with at least 10 connections ^70^ to or from the core song circuit bodyIDs (https://github.com/connectome-neuprint/neuprint-python). Cell-type-specific connectivity matrices were generated based on a cell-typing priority of our cell type (Sup. Table 1, which may include multiple MANC v1.0 systematic cell types) followed by MANC v1.0 systematic type. These connectivity matrices were used to analyze connectivity among the tested neurons, the identified wing motor neurons ^41^, and the MANC neurons that connect to the core song circuit neurons. Core song circuit neurons had >10% of their input/output to at least two other core neurons. This metric was used to identify neurons in the MANC with core-like features (Sup. Figure 14D). Circuit diagrams were generated in Cytoscape (https://cytoscape.org/) ^87^. Custom R scripts were used for all circuit structure analyses and for generating all non-circuit diagram circuit structure figures.

## Figures#

Details on figure panel creation are described in the appropriate section above. All figure panels were assembled in Adobe Illustrator.

## References

1. Ewing, A. W. & Bennet-Clark, H. C. The Courtship Songs of Drosophila. Behaviour 31, 288–301 (1968).

2. Spieth, H. T. Courtship Behavior in *Drosophila*. Annu. Rev. Entomol. 19, 385–405 (1974).

3. Arthur, B. J., Sunayama-Morita, T., Coen, P., Murthy, M. & Stern, D. L. Multi-channel acoustic recording and automated analysis of Drosophila courtship songs. BMC Biol. 11, 11 (2013).

4. Cowling, D. E. & Burnet, B. Courtship songs and genetic control of their acoustic characteristics in sibling species of the Drosophila melanogaster subgroup. Anim. Behav. 29, 924–935 (1981).

5. Siegel, R. W. & Hall, J. C. Conditioned responses in courtship behavior of normal and mutant Drosophila. Proc. Natl. Acad. Sci. 76, 3430–3434 (1979).

6. Zhang, S. X., Rogulja, D. & Crickmore, M. A. Recurrent Circuitry Sustains Drosophila Courtship Drive While Priming Itself for Satiety. Curr. Biol. 29, 3216–3228.e9 (2019).

7. Lin, H. H. et al. A nutrient-specific gut hormone arbitrates between courtship and feeding. Nature 602, 632–638 (2022).

8. Coen, P. et al. Dynamic sensory cues shape song structure in Drosophila. Nature 507, 233–237 (2014).

9. Clemens, J. et al. Discovery of a New Song Mode in Drosophila Reveals Hidden Structure in the Sensory and Neural Drivers of Behavior. Curr. Biol. 28, 2400–2412.e6 (2018).

10. Coen, P., Xie, M., Clemens, J. & Murthy, M. Sensorimotor Transformations Underlying Variability in Song Intensity during Drosophila Courtship. Neuron 89, 629–644 (2016).

11. Clemens, J. et al. Connecting Neural Codes with Behavior in the Auditory System of Drosophila. Neuron (2015). doi:10.1016/j.neuron.2015.08.014

12. Deutsch, D., Clemens, J., Thiberge, S. Y., Guan, G. & Murthy, M. Shared Song Detector Neurons in Drosophila Male and Female Brains Drive Sex-Specific Behaviors. Curr. Biol. 29, 3200–3215.e5 (2019).

13. Wang, K. et al. Neural circuit mechanisms of sexual receptivity in Drosophila females. Nat. 2020 5897843 589, 577–581 (2020).

14. Baker, C. A. et al. Neural network organization for courtship-song feature detection in Drosophila. Curr. Biol. 32, 3317–3333.e7 (2022).

15. Stockinger, P., Kvitsiani, D., Rotkopf, S., Tirián, L. & Dickson, B. J. Neural circuitry that governs Drosophila male courtship behavior. Cell 121, 795–807 (2005).

16. Lee, G. et al. Spatial, Temporal, and Sexually Dimorphic Expression Patterns of the fruitless Gene in the Drosophila Central Nervous System. J Neurobiol 43, 404–426 (2000).

17. Lee, G., Hall, J. C. & Park, J. H. DOUBLESEX GENE EXPRESSION IN THE CENTRAL NERVOUS SYSTEM OF DROSOPHILA MELANOGASTER. 10.1080/01677060216292 16, 229–248 (2009).

18. Manoli, D. S. et al. Male-specific fruitless specifies the neural substrates of Drosophila courtship behaviour. Nature 436, 395–400 (2005).

19. Rideout, E. J., Dornan, A. J., Neville, M. C., Eadie, S. & Goodwin, S. F. Control of sexual differentiation and behavior by the doublesex gene in Drosophila melanogaster. Nat. Neurosci. 2010 134 13, 458–466 (2010).

20. Bates, A. S. et al. Complete Connectomic Reconstruction of Olfactory Projection Neurons in the Fly Brain. Curr. Biol. 30, 3183–3199.e6 (2020).

21. Clowney, E. J., Iguchi, S., Bussell, J. J., Scheer, E. & Ruta, V. Multimodal Chemosensory Circuits Controlling Male Courtship in Drosophila. Neuron 87, 1036–1049 (2015).

22. Jefferis, G. S. X. E. et al. Comprehensive maps of Drosophila higher olfactory centers: spatially segregated fruit and pheromone representation. Cell 128, 1187–203 (2007).

23. Kallman, B. R., Kim, H. & Scott, K. Excitation and inhibition onto central courtship neurons biases drosophila mate choice. Elife 4, (2015).

24. Kohl, J., Ostrovsky, A. D., Frechter, S. & Jefferis, G. S. X. E. A Bidirectional Circuit Switch Reroutes Pheromone Signals in Male and Female Brains. Cell 155, 1610 (2013).

25. Kurtovic, A., Widmer, A. & Dickson, B. J. A single class of olfactory neurons mediates behavioural responses to a Drosophila sex pheromone. Nat. 2006 4467135 446, 542–546 (2007).

26. Marin, E. C., Jefferis, G. S. X. ., Komiyama, T., Zhu, H. & Luo, L. Representation of the Glomerular Olfactory Map in the Drosophila Brain. Cell 109, 243–255 (2002).

27. Ribeiro, I. M. A. et al. Visual Projection Neurons Mediating Directed Courtship in Drosophila. Cell 174, 607–621.e18 (2018).

28. Wong, A. M., Wang, J. W. & Axel, R. Spatial Representation of the Glomerular Map in the Drosophila Protocerebrum. Cell 109, 229–241 (2002).

29. Hindmarsh Sten, T., Li, R., Otopalik, A. & Ruta, V. Sexual arousal gates visual processing during Drosophila courtship. Nature 595, 549–553 (2021).

30. Lillvis, J. L. et al. Rapid reconstruction of neural circuits using tissue expansion and light sheet microscopy. Elife 11, (2022).

31. von Philipsborn, A. C. et al. Neuronal control of Drosophila courtship song. Neuron 69, 509–22 (2011).

32. Kohatsu, S., Koganezawa, M. & Yamamoto, D. Female contact activates male-specific interneurons that trigger stereotypic courtship behavior in Drosophila. Neuron 69, 498–508 (2011).

33. Kohatsu, S. & Yamamoto, D. Visually induced initiation of Drosophila innate courtship-like following pursuit is mediated by central excitatory state. Nat. Commun. 6, 1–9 (2015).

34. Shirangi, T. R. R., Stern, D. L. L. & Truman, J. W. W. Motor Control of Drosophila Courtship Song. Cell Rep. 5, 678–86 (2013).

35. Shirangi, T. R., Wong, A. M., Truman, J. W. & Stern, D. L. Doublesex Regulates the Connectivity of a Neural Circuit Controlling Drosophila Male Courtship Song. Dev. Cell 37, 533–544 (2016).

36. Ewing, A. W. The Neuromuscular Basis of Courtship song in Drosophila: The Role of the Indirect Flight Muscles. J. Comp. Physiol 119, 249–265 (1977).

37. Ewing, A. W. The Neuromuscular Basis of Courtship Song in Drosophila: The Role of the Direct and Axillary Wing Muscles. J. Comp. Physiol 130, 87–93 (1979).

38. Ewing, A. W. The role of feedback during singing and flight in Drosophila melanogaster. Physiol. Entomol. 4, 329–337 (1979).

39. O’Sullivan, A. et al. Multifunctional Wing Motor Control of Song and Flight. Curr. Biol. 28, 2705–2717.e4 (2018).

40. Marin, E. C. et al. Systematic annotation of a complete adult male Drosophila nerve cord connectome reveals principles of functional organisation. bioRxiv 2023.06.05.543407 (2023). doi:10.1101/2023.06.05.543407

41. Cheong, H. S. J. et al. Transforming descending input into behavior: The organization of premotor circuits in the Drosophila Male Adult Nerve Cord connectome. bioRxiv 2023.06.07.543976 (2023). doi:10.1101/2023.06.07.543976

42. Takemura, S.-Y. et al. A Connectome of the Male Drosophila Ventral Nerve Cord. bioRxiv 2023.06.05.543757 (2023). doi:10.1101/2023.06.05.543757

43. Cachero, S., Ostrovsky, A. D., Yu, J. Y., Dickson, B. J. & Jefferis, G. S. X. E. Sexual Dimorphism in the Fly Brain. Curr. Biol. 20, 1589–1601 (2010).

44. Yu, J. Y., Kanai, M. I., Demir, E., Jefferis, G. S. X. E. & Dickson, B. J. Cellular organization of the neural circuit that drives Drosophila courtship behavior. Curr. Biol. 20, 1602–14 (2010).

45. Mohammad, F. et al. Optogenetic inhibition of behavior with anion channelrhodopsins. Nat. Methods 14, 271–274 (2017).

46. Klapoetke, N. C. et al. Independent optical excitation of distinct neural populations. Nat. Methods 11, 338–346 (2014).

47. Arthur, B. J., et al. SongExplorer: A deep learning workflow for discovery and segmentation of animal acoustic communication signals. bioRxiv 2021.03.26.437280 (2021). doi:10.1101/2021.03.26.437280

48. Shiozaki, H. M., et al. Neural coding of distinct motor patterns during Drosophila courtship song. bioRxiv (2022). doi:10.1101/2022.12.14.520499

49. Nern, A., Pfeiffer, B. D. & Rubin, G. M. Optimized tools for multicolor stochastic labeling reveal diverse stereotyped cell arrangements in the fly visual system. 112, (2015).

50. Ding, Y. et al. Neural Evolution of Context-Dependent Fly Song. Curr. Biol. 29, 1089–1099.e7 (2019).

51. Calhoun, A. J., Pillow, J. W. & Murthy, M. Unsupervised identification of the internal states that shape natural behavior. Nat. Neurosci. 2019 2212 22, 2040–2049 (2019).

52. Bennet-Clark, H. C. & Ewing, A. W. Pulse interval as a critical parameter in the courtship song of Drosophila melanogaster. Anim. Behav. 17, 755–759 (1969).

53. Schilcher, F. von. The function of pulse song and sine song in the courtship of Drosophila melanogaster. Anim. Behav. 24, 622–625 (1976).

54. Dickinson, M. H. & Tu, M. S. The Function of Dipteran Flight Muscle. Comp. Biochem. Physiol. Part A Physiol. 116, 223–238 (1997).

55. Trimarchi, J. R. & Schneiderman, A. M. The motor neurons innervating the direct flight muscles of Drosophila melanogaster are morphologically specialized. J. Comp. Neurol. 340, 427–443 (1994).

56. Ehrhardt, E. et al. Single-cell type analysis of wing premotor circuits in the ventral nerve cord of Drosophila melanogaster. bioRxiv 2023.05.31.542897 (2023). doi:10.1101/2023.05.31.542897

57. Marder, E. & Bucher, D. Understanding Circuit Dynamics Using the Stomatogastric Nervous System of Lobsters and Crabs. 10.1146/annurev.physiol.69.031905.161516 69, 291–316 (2007).

58. Peña, F., Parkis, M. A., Tryba, A. K. & Ramirez, J. M. Differential contribution of pacemaker properties to the generation of respiratory rhythms during normoxia and hypoxia. Neuron 43, 105–117 (2004).

59. Lieske, S. P., Thoby-Brisson, M., Telgkamp, P. & Ramirez, J. M. Reconfiguration of the neural network controlling multiple breathing patterns: eupnea, sighs and gasps. Nat. Neurosci. 3, 600–607 (2000).

60. Briggman, K. L. & Kristan, W. B. Imaging Dedicated and Multifunctional Neural Circuits Generating Distinct Behaviors. J. Neurosci. 26, 10925–10933 (2006).

61. Esch, T., Mesce, K. A. & Kristan, W. B. Evidence for sequential decision making in the medicinal leech. J. Neurosci. 22, 11045–11054 (2002).

62. Tomaru, M. & Yamada, H. Courtship of Drosophila, with a special interest in courtship songs. Low Temp. Sci. 69, 61–85 (2011).

63. Tracy, C. B., Nguyen, J., Abraham, R. & Shirangi, T. R. Evolution of sexual size dimorphism in the wing musculature of Drosophila. PeerJ **2020**, e8360 (2020).

64. Briggman, K. L. & Kristan, W. B. Multifunctional pattern-generating circuits. Annu. Rev. Neurosci. 31, 271–294 (2008).

65. Xu, C. S. et al. Enhanced FIB-SEM systems for large-volume 3D imaging. Elife 6, (2017).

66. Sheridan, A. et al. Local shape descriptors for neuron segmentation. Nat. Methods 2022 202 20, 295–303 (2022).

67. Schlegel, P. et al. A consensus cell type atlas from multiple connectomes reveals principles of circuit stereotypy and variation. bioRxiv 2023.06.27.546055 (2023). doi:10.1101/2023.06.27.546055

68. Lesser, E. et al. Synaptic architecture of leg and wing motor control networks in Drosophila. bioRxiv 2023.05.30.542725 (2023). doi:10.1101/2023.05.30.542725

69. Phelps, J. S. et al. Reconstruction of motor control circuits in adult Drosophila using automated transmission electron microscopy. Cell 184, 759–774.e18 (2021).

70. Schlegel, P. et al. Whole-brain annotation and multi-connectome cell typing quantifies circuit stereotypy in Drosophila. bioRxiv 2023.06.27.546055 (2023). doi:10.1101/2023.06.27.546055

71. Dorkenwald, S. et al. Neuronal wiring diagram of an adult brain. bioRxiv (2023). doi:10.1101/2023.06.27.546656

72. Scheffer, L. K. et al. A connectome and analysis of the adult drosophila central brain. Elife 9, 1–74 (2020).

73. Phelps, J. S. et al. Reconstruction of motor control circuits in adult Drosophila using automated transmission electron microscopy. Cell (2021). doi:10.1016/j.cell.2020.12.013

74. Dionne, H., Hibbard, K. L., Cavallaro, A., Kao, J.-C. & Rubin, G. M. Genetic Reagents for Making Split-GAL4 Lines in Drosophila. Genetics 209, 31–35 (2018).

75. Tirian, L. & Dickson, B. The VT GAL4, LexA, and split-GAL4 driver line collections for targeted expression in the Drosophila nervous system. bioRxiv 198648 (2017). doi:10.1101/198648

76. Otsuna, H., Ito, M. & Kawase, T. Color depth MIP mask search: a new tool to expedite Split-GAL4 creation. bioRxiv 318006 (2018). doi:10.1101/318006

77. Wu, M. et al. Visual projection neurons in the Drosophila lobula link feature detection to distinct behavioral programs. Elife 5, (2016).

78. Mellert, D. J., Knapp, J. M., Manoli, D. S., Meissner, G. W. & Baker, B. S. Midline crossing by gustatory receptor neuron axons is regulated by fruitless, doublesex and the Roundabout receptors. Development 137, 323–332 (2010).

79. Wang, Y. et al. EASI-FISH for thick tissue defines lateral hypothalamus spatio-molecular organization. Cell 184, 6361–6377.e24 (2021).

80. Eddison, M. & Ihrke, G. Expansion-assisted iterative fluorescence in situ hybridization (EASI-FISH) in Drosophila CNS. Protocols.IO (2022). Available at: https://www.protocols.io/view/expansion-assisted-iterative-fluorescence-in-situ-cg7ktzkw.html. (Accessed: 17th July 2023)

81. Meissner, G. W. et al. Mapping Neurotransmitter Identity in the Whole-Mount Drosophila Brain Using Multiplex High-Throughput Fluorescence in Situ Hybridization. Genetics genetics.301749.2018 (2018). doi:10.1534/genetics.118.301749

82. Long, X., Colonell, J., Wong, A. M., Singer, R. H. & Lionnet, T. Quantitative mRNA imaging throughout the entire Drosophila brain. Nat. Methods 14, 703–706 (2017).

83. Bogovic, J. A. et al. An unbiased template of the Drosophila brain and ventral nerve cord. PLoS One 15, e0236495 (2020).

84. Wan, Y., Otsuna, H., Chien, C.-B. & Hansen, C. FluoRender: An Application of 2D Image Space Methods for 3D and 4D Confocal Microscopy Data Visualization in Neurobiology Research. IEEE Pacific Vis. Symp. 201–208 (2012).

85. Dana, H. et al. High-performance calcium sensors for imaging activity in neuronal populations and microcompartments. Nat. Methods 16, 649–657 (2019).

86. Clements, J. et al. NeuronBridge: an intuitive web application for neuronal morphology search across large data sets. bioRxiv 2022.07.20.500311 (2022). doi:10.1101/2022.07.20.500311

87. Shannon, P. et al. Cytoscape: A Software Environment for Integrated Models of Biomolecular Interaction Networks. Genome Res. 13, 2498–2504 (2003).

