## Supplemental Figures for "Nested neural circuits generate distinct acoustic signals during Drosophila courtship"

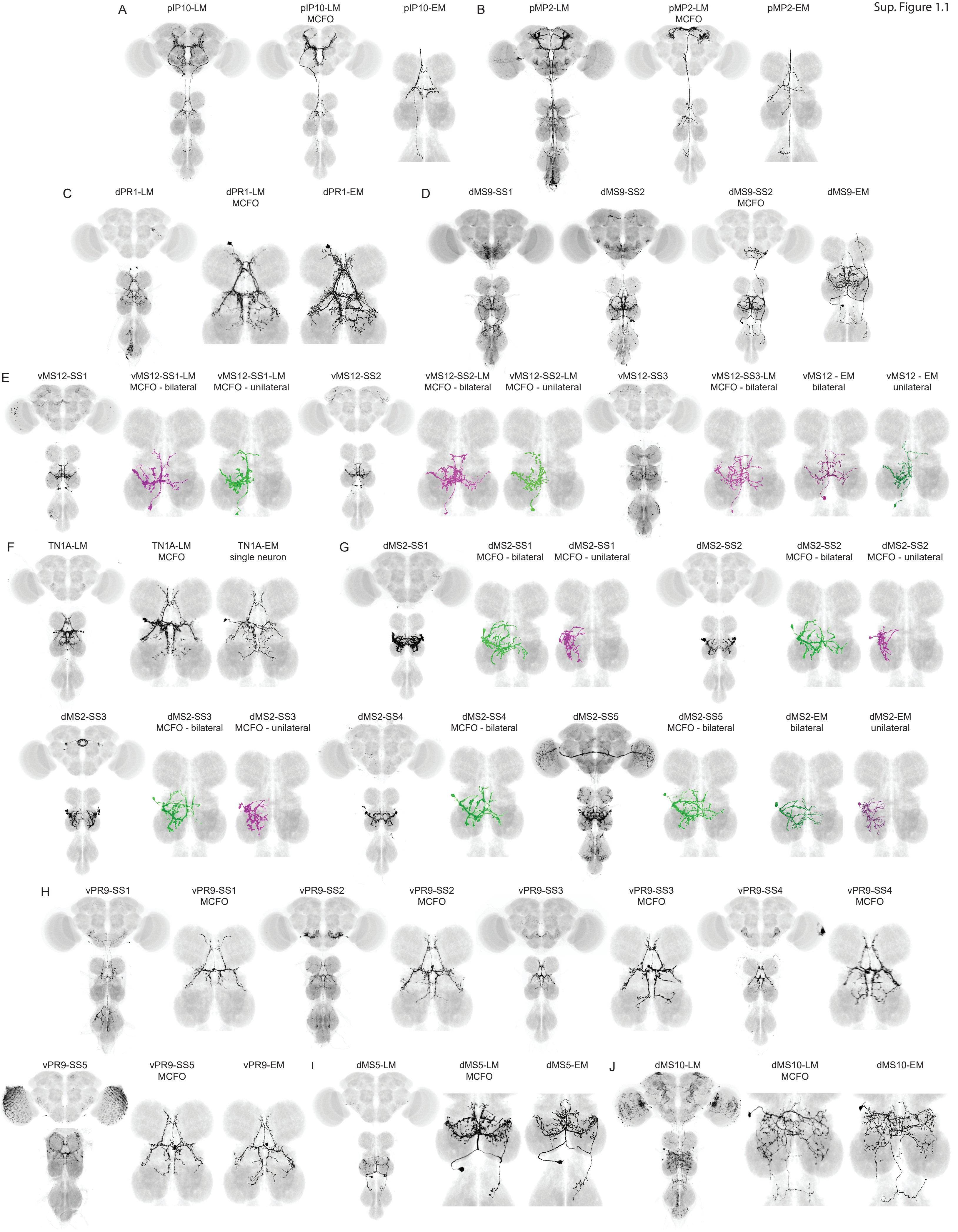


Sup. Figure 1.1

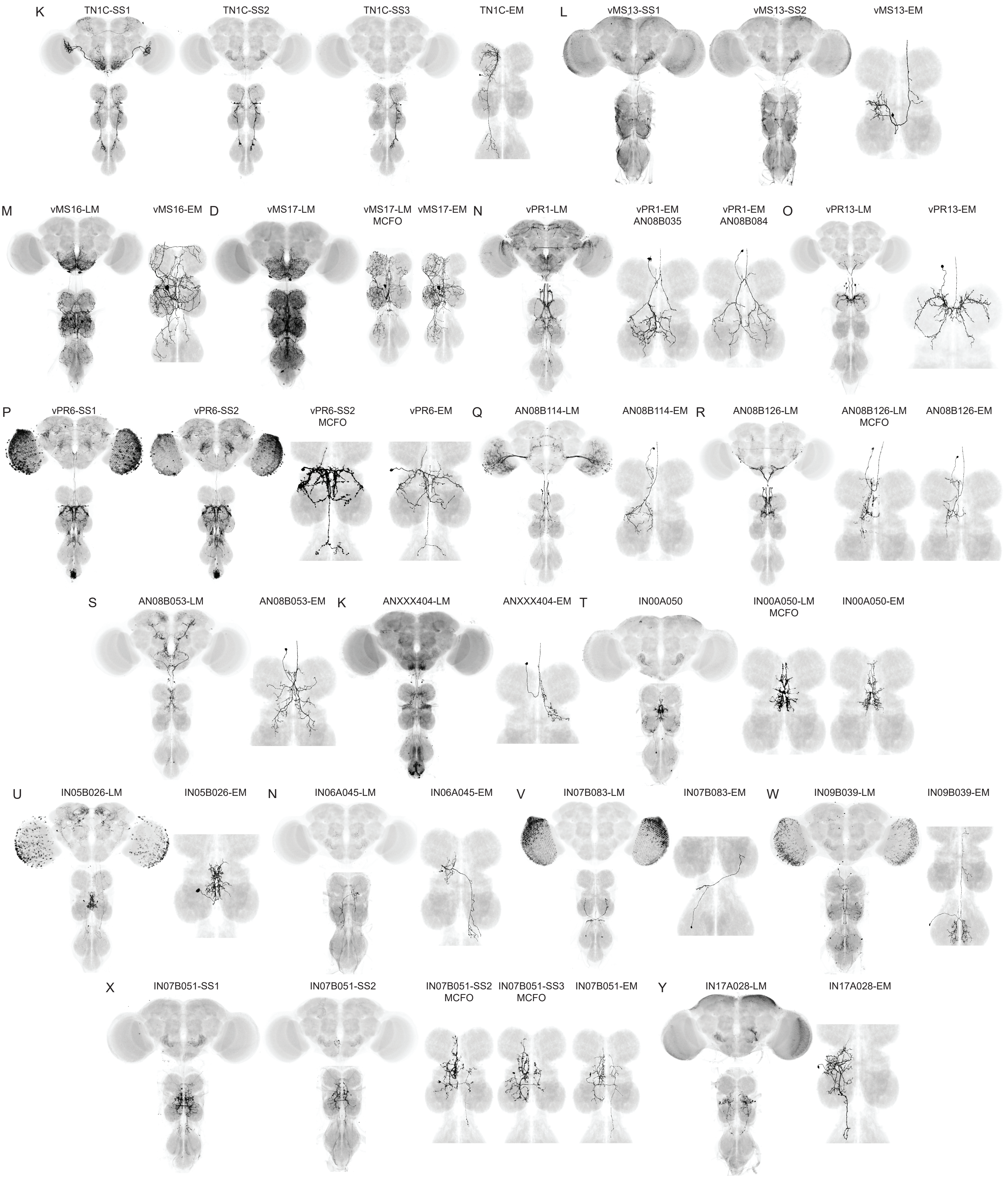
K
TN1C-SS1
TN1C-SS2
TN1C-SS3
TN1C-EM
L
vMS13-SS1
vMS13-SS2
vMS13-EM
M
vMS16-LM
vMS16-EM
D
vMS17-LM
vMS17-LM MCFO
vMS17-EM
N
vPR1-LM
vPR1-EM AN08B035
vPR1-EM AN08B084
O
vPR13-LM
vPR13-EM
P
vPR6-SS1
vPR6-SS2
vPR6-SS2 MCFO
vPR6-EM
Q
AN08B114-LM
AN08B114-EM
R
AN08B126-LM
AN08B126-LM MCFO
AN08B126-EM
S
AN08B053-LM
AN08B053-EM
K
ANXXX404-LM
ANXXX404-EM
T
IN00A050
IN00A050-LM MCFO
IN00A050-EM
U
IN05B026-LM
IN05B026-EM
N
IN06A045-LM
IN06A045-EM
V
IN07B083-LM
IN07B083-EM
W
IN09B039-LM
IN09B039-EM
X
IN07B051-SS1
IN07B051-SS2
IN07B051-SS2 MCFO
IN07B051-SS3 MCFO
IN07B051-EM
Y
IN17A028-LM
IN17A028-EM

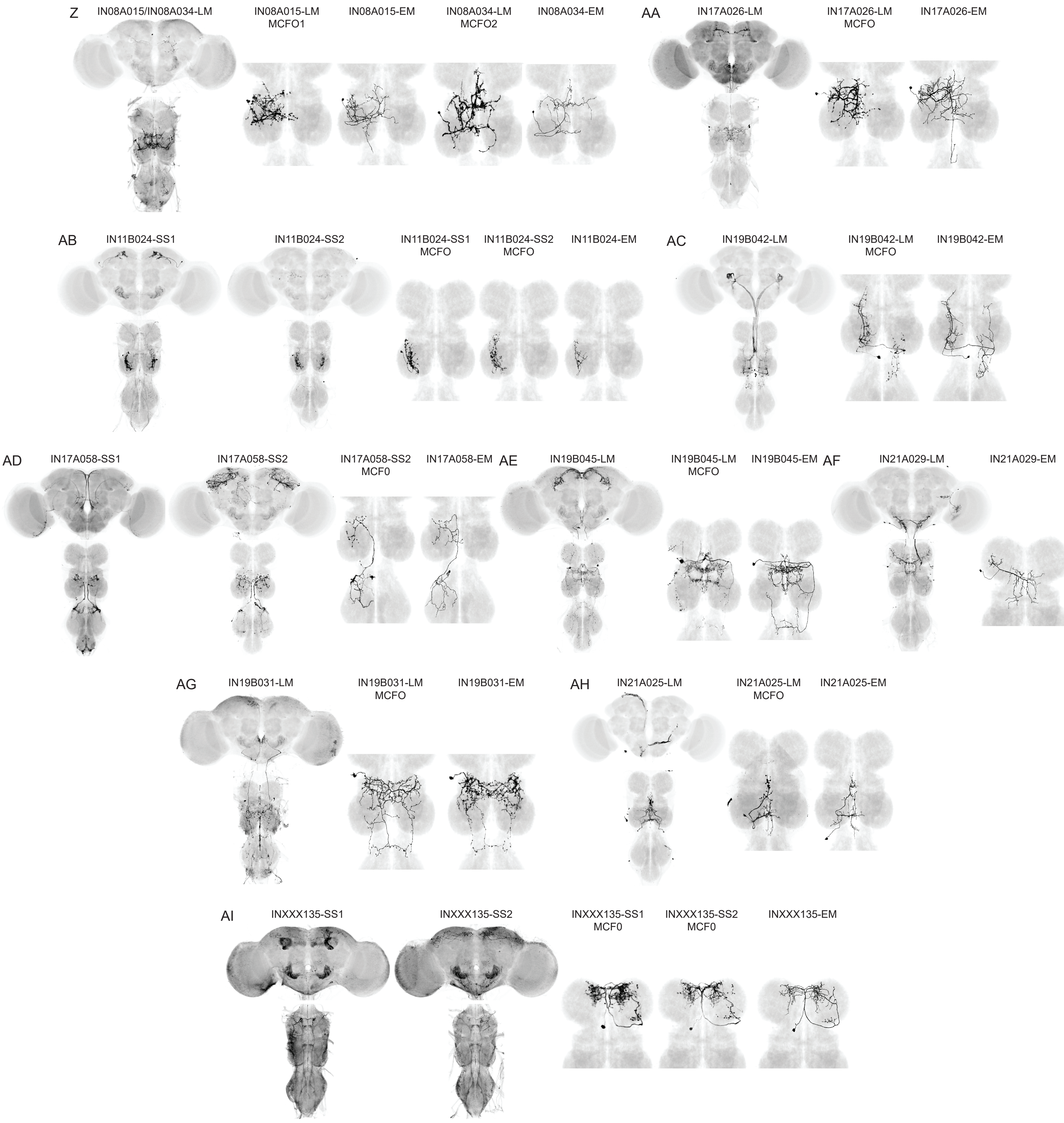


Sup. Figure 1 Representative example images of split-GAL4 lines, individual stochastically labeled neurons from split-GAL4 lines (where available), and neuron match candidates in the MANC volume. LM=light microscopy, SS=stable split, MCFO=multi-color flip-out, EM=electron microscopy. See Table 1 and methods for details on each split-GAL4 line and EM matching. Confocal stacks of every split-GAL4 line are available at https://splitgal4.janelia.org/cgi-bin/splitgal4.cgi and 3D meshes of every EM body ID are available at neuprint.janelia.org.

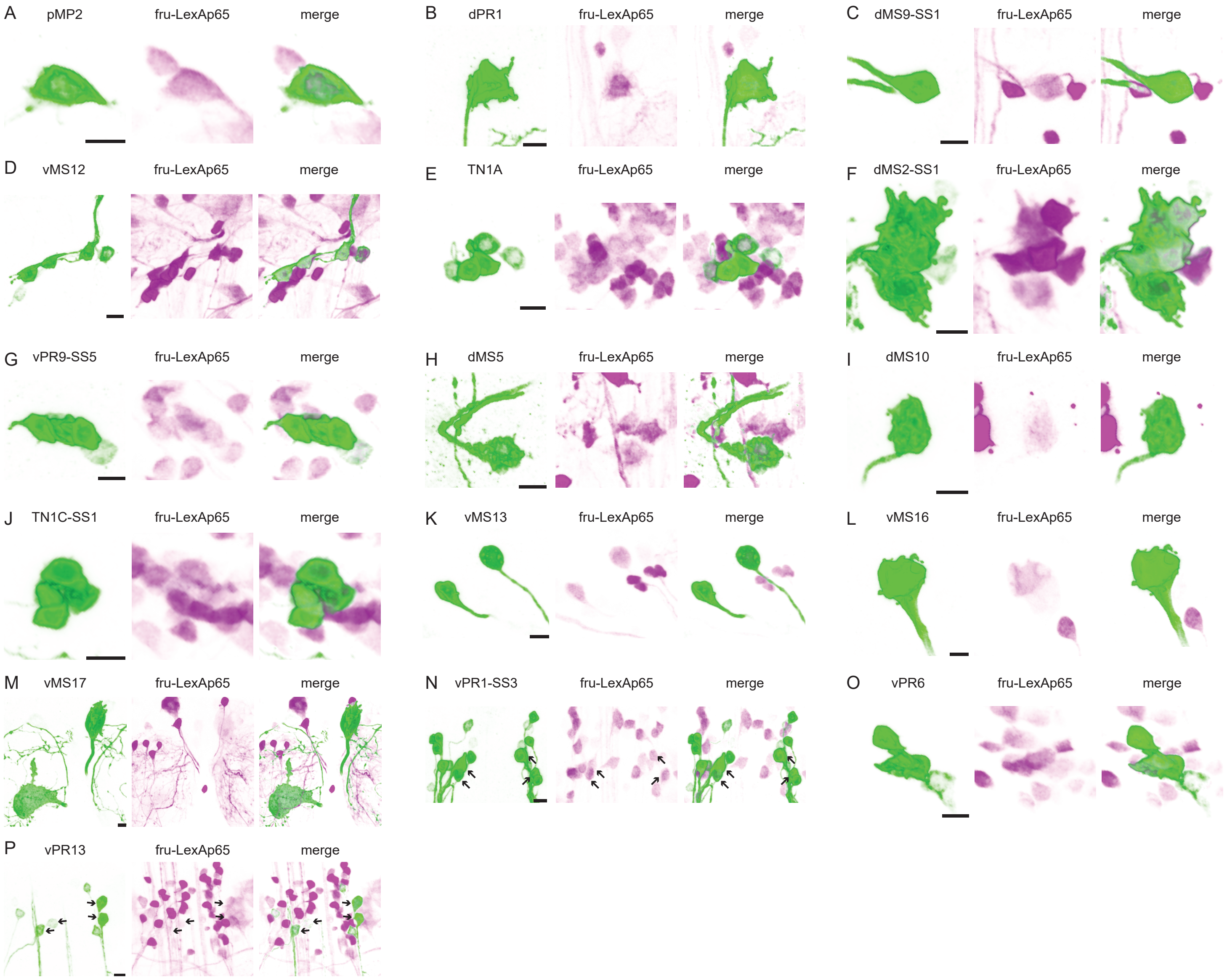


Sup. Figure 2 Identification of fru+ neurons via split-GAL4 (green) colocalization with fru-LexAp65 (magenta). Representative examples of each cell type shown. Arrows indicate fru+ neurons among a population of fru+ and fru- neurons. Scale bars = 5 μm. See Table 1 for details of each split-GAL4 line and quantification.

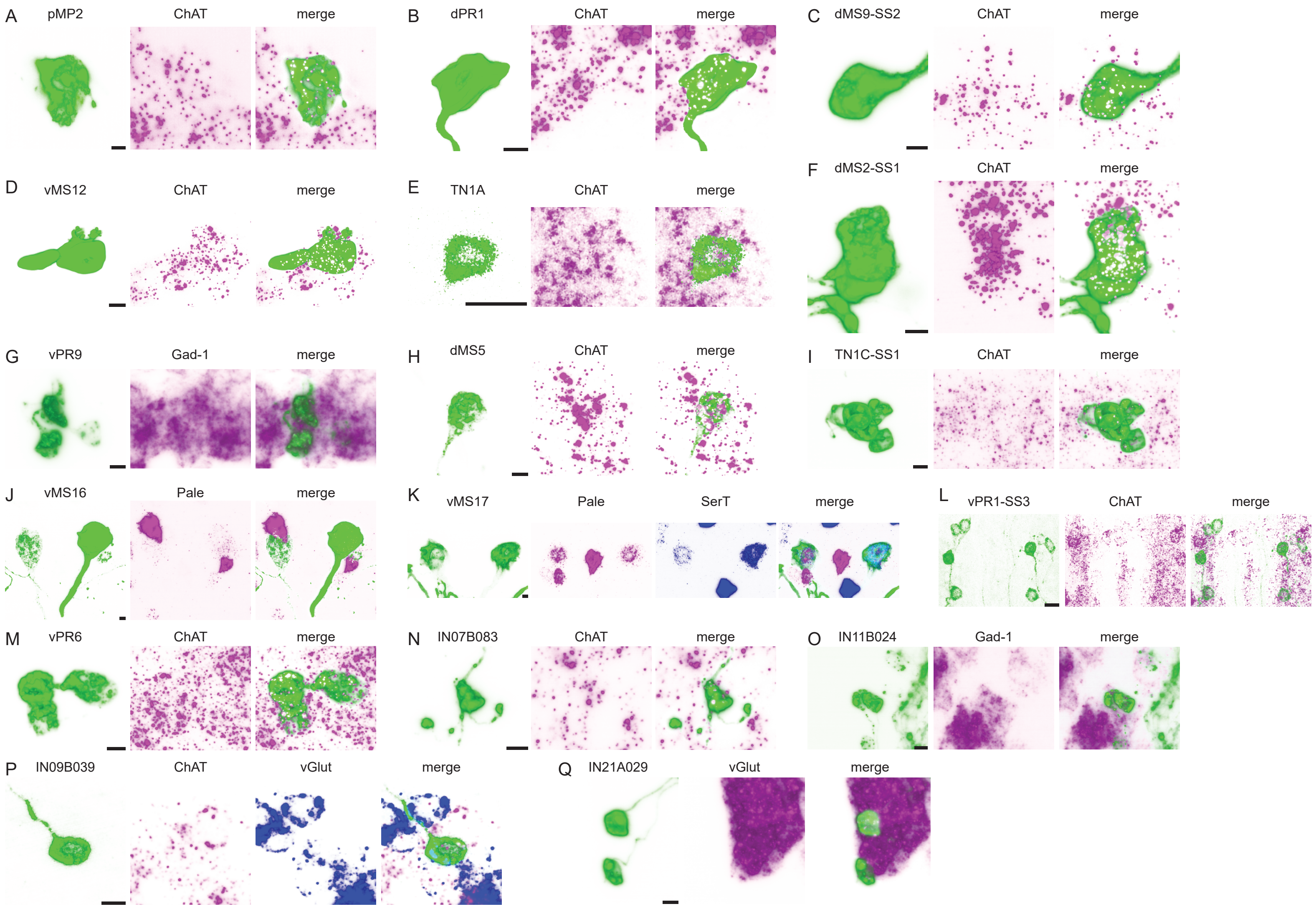


Sup. Figure 3 Neurotransmitter identification by EASI-FISH. Representative images of split-GAL4>GFP colocalization with probes targeting the indicated mRNA. ChAT = acetylcholine, Gad-1 = GABA, pale = dopamine, SerT = serotonin, vGlut = glutamate. Scale bars = 5 μm.

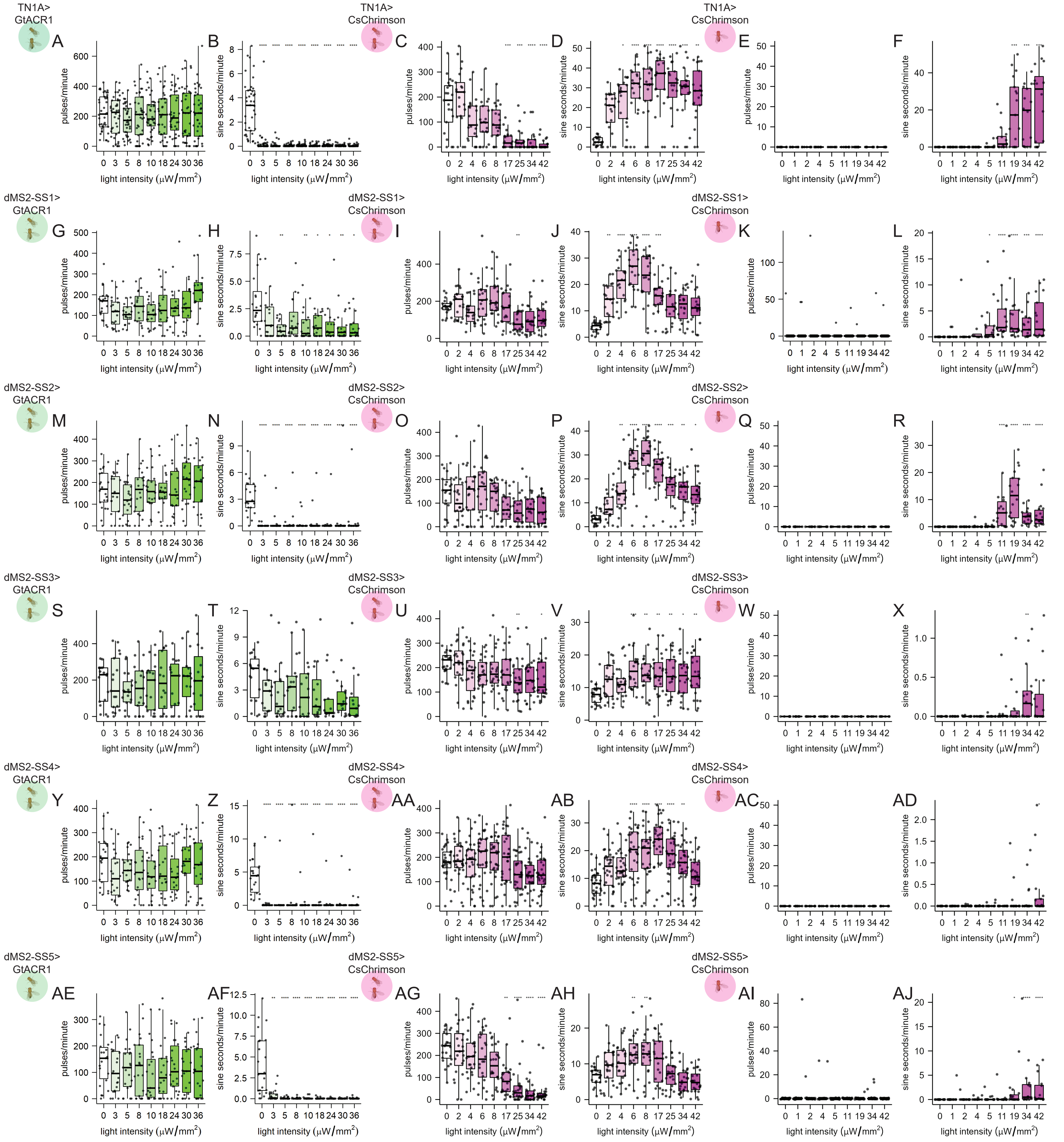


Sup. Figure 4 Quantification of pulses/minute or sine seconds/minute in courting split-GAL4>GtACR1, courting split-GAL4>CsChrimson, and isolated split-GAL4>CsChrimson males without light stimuli (0) and at increasing light stimulus intensities. split-GAL4, effector, and assay indicated to the left of each set of plots. p-values calculated via Kruskal-Wallis tests and Dunn's pairwise comparisons with Bonferroni correction. *p<0.05, **p<0.005, ***p<0.0005, ****p<0.00005.

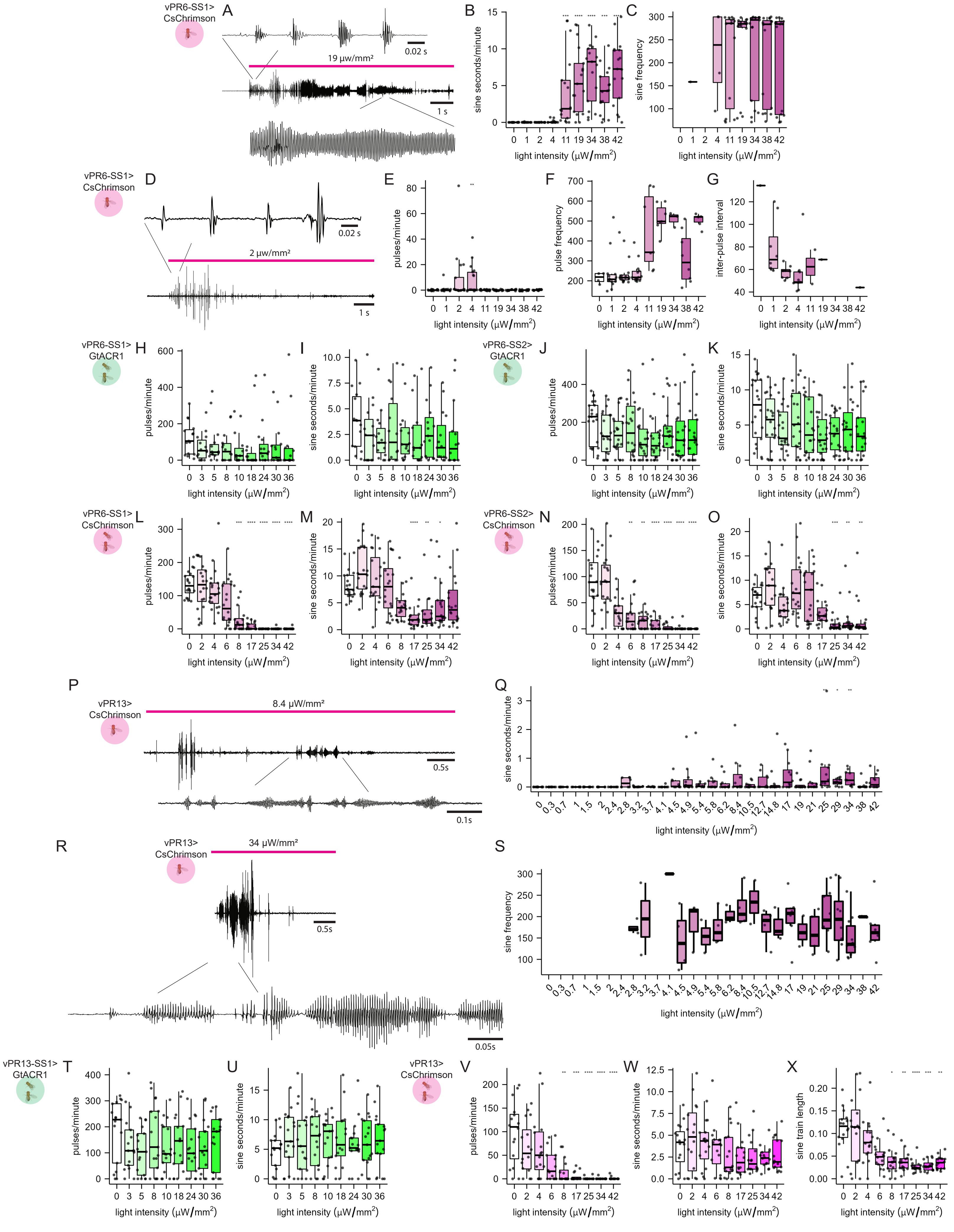

vPR6-SS1> CsChrimson
A
0.02 s
19 μw/mm²
1 s
B
sine seconds/minute
light intensity (μW/mm²)
C
sine frequency
light intensity (μW/mm²)
vPR6-SS1> CsChrimson
D
0.02 s
2 μw/mm²
1 s
E
pulses/minute
light intensity (μW/mm²)
F
pulse frequency
light intensity (μW/mm²)
G
inter-pulse interval
light intensity (μW/mm²)
vPR6-SS1> GtACR1
H
pulses/minute
light intensity (μW/mm²)
I
sine seconds/minute
light intensity (μW/mm²)
vPR6-SS2> GtACR1
J
pulses/minute
light intensity (μW/mm²)
K
sine seconds/minute
light intensity (μW/mm²)
vPR6-SS1> CsChrimson
L
pulses/minute
light intensity (μW/mm²)
M
sine seconds/minute
light intensity (μW/mm²)
vPR6-SS2> CsChrimson
N
pulses/minute
light intensity (μW/mm²)
O
sine seconds/minute
light intensity (μW/mm²)
P
vPR13> CsChrimson
8.4 μW/mm²
0.5s
0.1s
Q
sine seconds/minute
light intensity (μW/mm²)
R
vPR13> CsChrimson
34 μW/mm²
0.5s
0.05s
S
sine frequency
light intensity (μW/mm²)
vPR13-SS1> GtACR1
T
pulses/minute
light intensity (μW/mm²)
U
sine seconds/minute
light intensity (μW/mm²)
vPR13> CsChrimson
V
pulses/minute
light intensity (μW/mm²)
W
sine seconds/minute
light intensity (μW/mm²)
X
sine train length
light intensity (μW/mm²)

Sup. Figure 5 A) Representative example of polycyclic pulse-like trains and sine-like trains produced in isolated vPR6>CsChrimson males by 19 μW/mm² red light stimuli. B) Quantification of sine seconds/minute and C) sine frequency in isolated vPR6>CsChrimson males without red light stimuli (0) and at increasing red light stimulus intensities. D) Representative example of pulse trains produced in isolated vPR6>CsChrimson males by 2 μW/mm² red light stimuli. E) Quantification of pulses/minute, F) pulse carrier frequency, and G) inter-pulse interval in isolated vPR6>CsChrimson males without red light stimuli (0) and at increasing red light stimulus intensities. H) Quantification of pulses/minute and I) sine seconds/minute in courting vPR6-SS1>GtACR1 males without green light stimuli (0) and at increasing green light stimulus intensities. J-K) Same as H-I, but for vPR6-SS2. L-M) Same as H-K but for vPR6>CsChrimson males and red light stimuli. P) Representative example of wing sounds, including sine trains, produced in isolated vPR13>CsChrimson males by 8.4 μW/mm² red light stimuli. Q) Quantification of sine seconds/minute in isolated vPR13>CsChrimson males without red light stimuli (0) and at increasing red light stimulus intensities. R) Representative example of wing sounds, including high amplitude sine-like trains, produced in isolated vPR13>CsChrimson males by 34 μW/mm² red light stimuli. S) Same as Q, but for sine carrier frequency. T) Quantification of pulses/minute, U) sine seconds/minute in courting vPR13>GtACR1 males without green light stimuli (0) and at increasing green light stimulus intensities. V-W) Same as T-U, but for vPR13>CsChrimson males and red light. X) Same as (W), but for sine train length. p-values calculated via Kruskal-Wallis tests and Dunn's pairwise comparisons with Bonferroni correction. *p<0.05, **p<0.005, ***p<0.0005, ****p<0.00005.

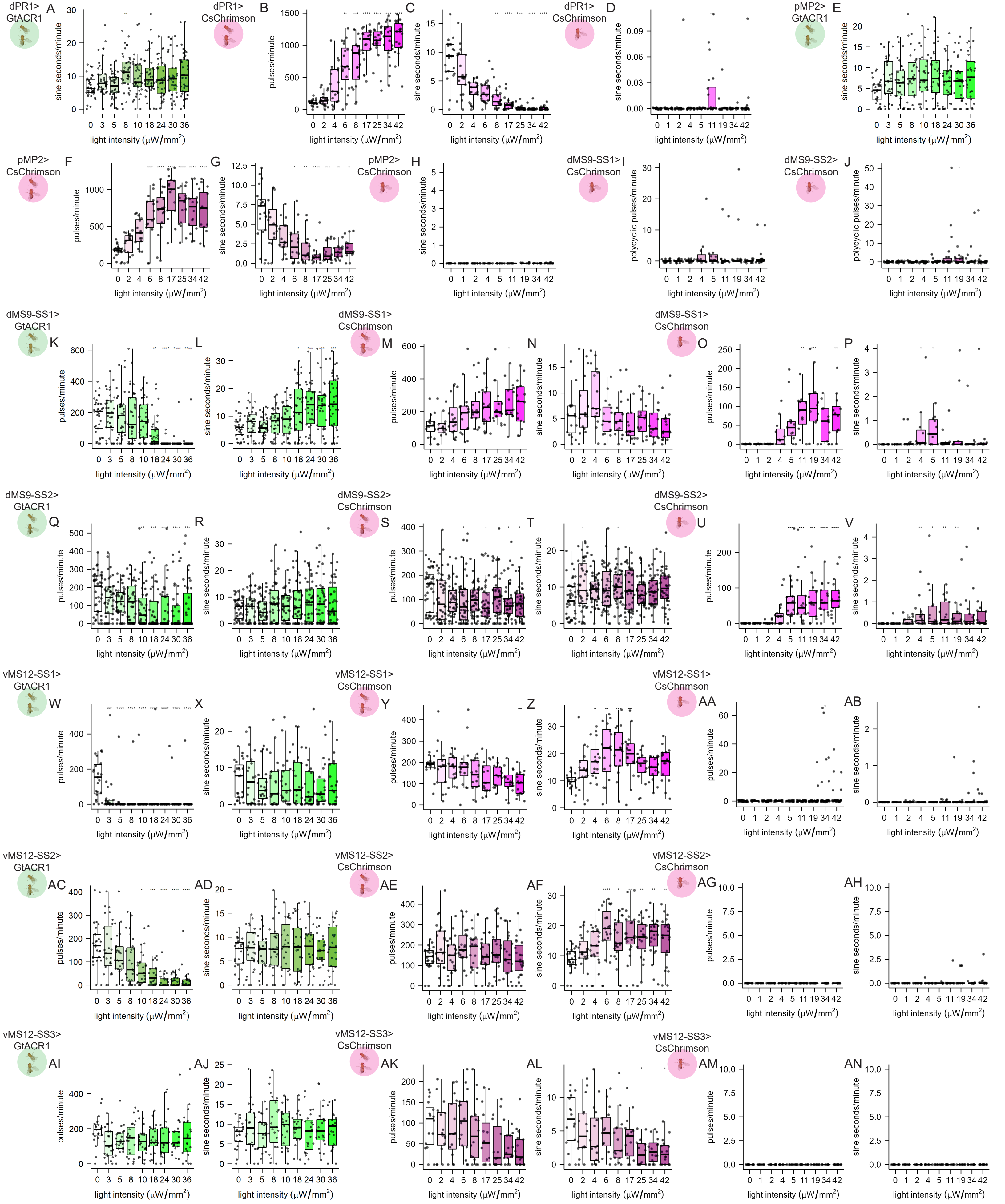

Sup. Figure 6 Quantification of pulses/minute or sine seconds/minute in courting split-GAL4>GtACR1, courting split-GAL4>CsChrimson, and isolated split-GAL4>CsChrimson males without light stimuli (0) and at increasing light stimulus intensities. Split-GAL4, effector, and assay indicated to the left of each set of plots.p-values calculated via Kruskal-Wallis tests and Dunn's pairwise comparisons with Bonferroni correction. *p<0.05, **p<0.005, ***p<0.0005, ****p<0.00005.

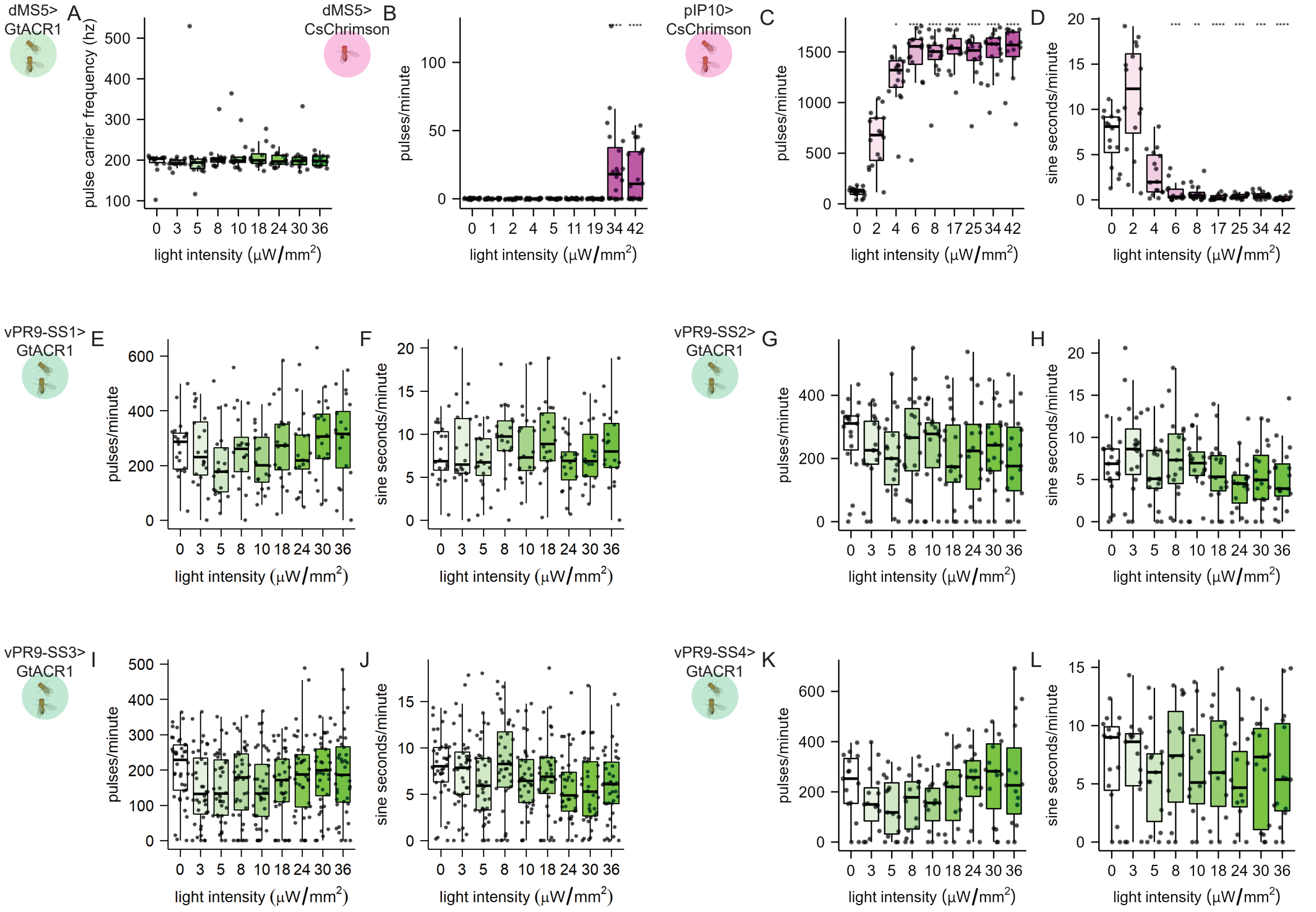


Sup. Figure 7 Quantification of pulses/minute or sine seconds/minute in courting split-GAL4>GtACR1 and isolated split-GAL4>CsChrimson males without light stimuli (0) and at increasing light stimulus intensities. split-GAL4, effector, and assay indicated to the left of each set of plots. p-values calculated via Kruskal-Wallis tests and Dunn's pairwise comparisons with Bonferroni correction. *p<0.05, **p<0.005, ***p<0.0005, ****p<0.00005.

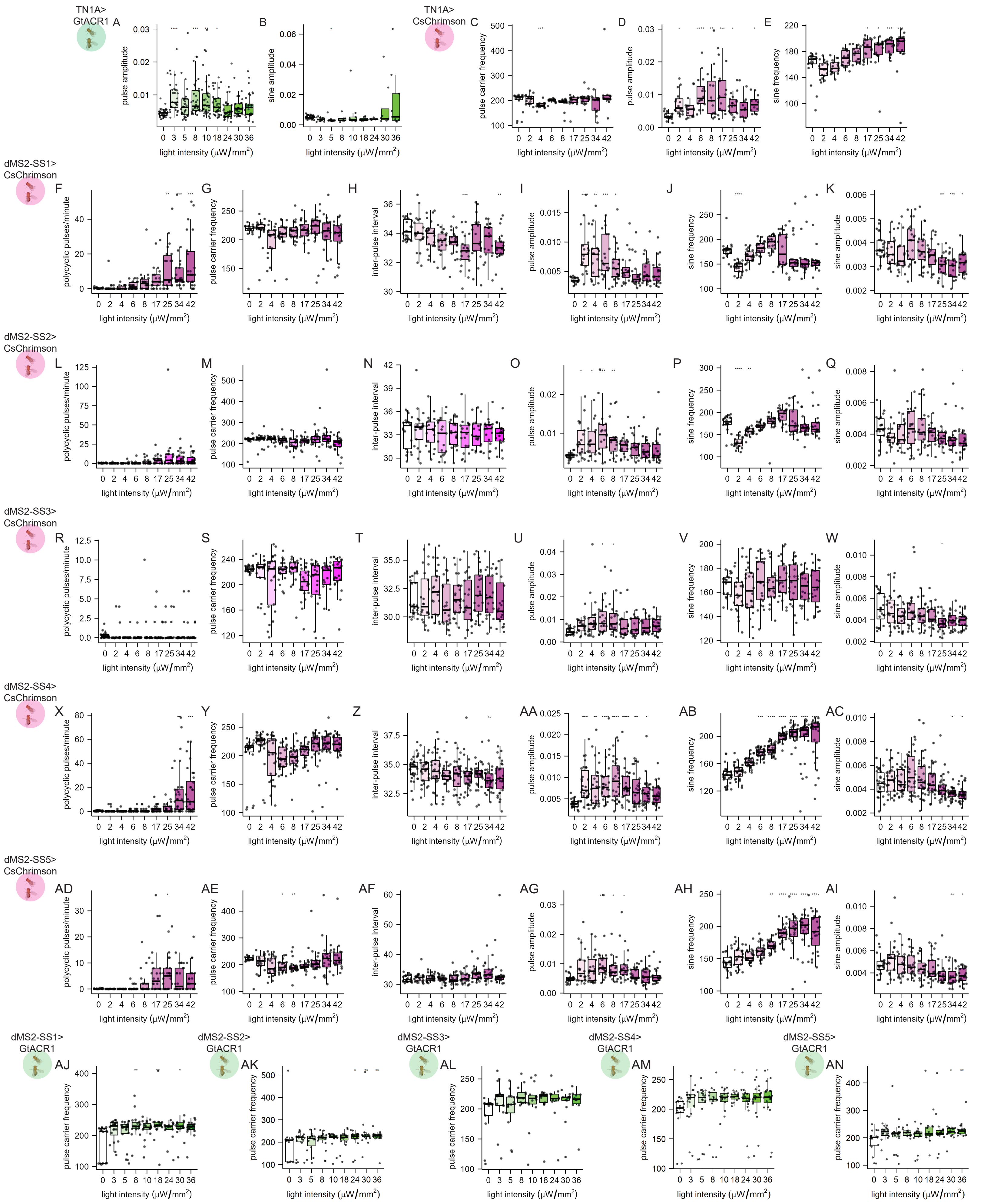

Sup. Figure 8 Quantification of pulse carrier frequency (hz), sine frequency (hz), pulse and sine amplutide (relative units), inter-pulse interval (ms), or polycyclic pulses/minute in courting split-GAL4>CsChrimson or split-GAL4>GtACR1 males without light stimuli (0) and increasing light stimulus intensities. split-GAL4 driver/neuron type, effector, and assay indicated to the left of each set of plots. p-values calculated via Kruskal-Wallis tests and Dunn's pairwise comparisons with Bonferroni correction. *p<0.05, **p<0.005, ***p<0.0005, ****p<0.00005.

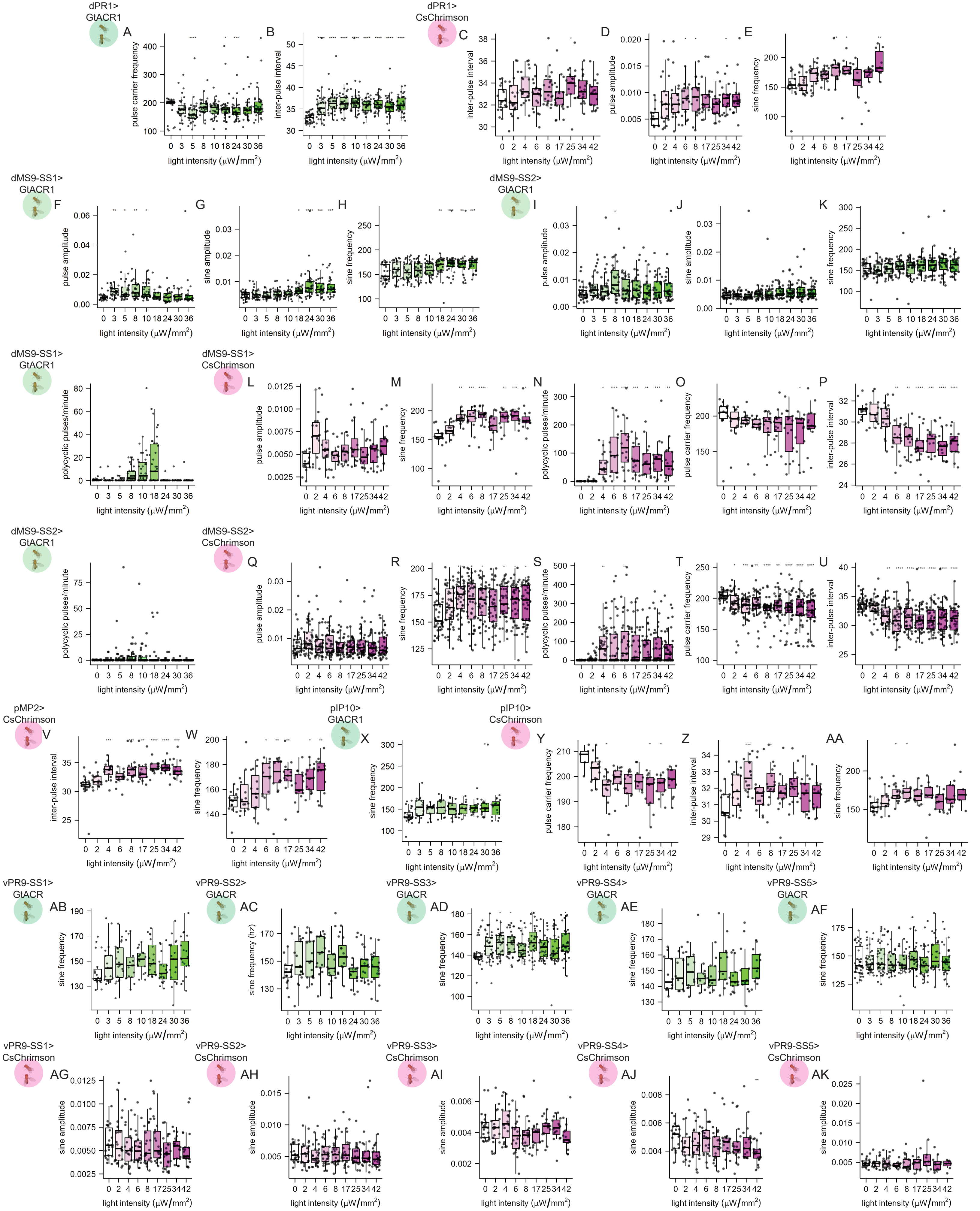


Sup. Figure 9 Quantification of pulse carrier frequency (hz), sine frequency (hz), pulse and sine amplitude (relative units), inter-pulse interval (ms), or polycyclic pulses/minute in courting split-GAL4>CsChrimson or split-GAL4>GtACR1 males without light stimuli (0) and increasing light stimulus intensities. split-GAL4 driver/neuron type, effector, and assay indicated to the left of each set of plots. p-values calculated via Kruskal-Wallis tests and Dunn's pairwise comparisons with Bonferroni correction. *p<0.05, **p<0.005, ***p<0.0005, ****p<0.00005.

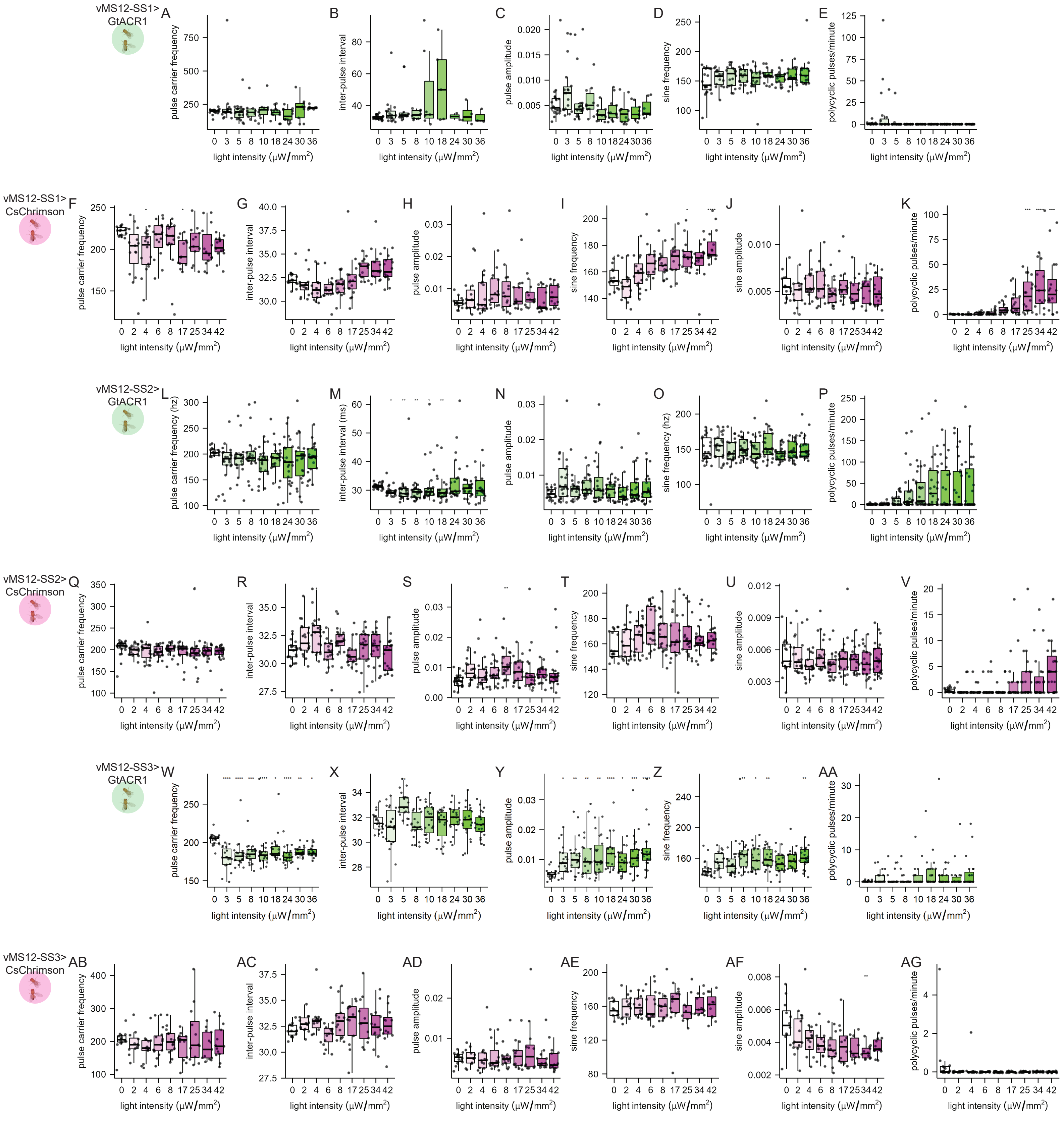


Sup. Figure 10 Quantification of pulse carrier frequency (hz), sine frequency (hz), pulse and sine amplutide (relative units), inter-pulse interval (ms), or polycyclic pulses/minute in courting split-GAL4>CsChrimson or split-GAL4>GtACR1 males without light stimuli (0) and increasing light stimulus intensities. split-GAL4 driver/neuron type, effector, and assay indicated to the left of each set of plots. p-values calculated via Kruskal-Wallis tests and Dunn's pairwise comparisons with Bonferroni correction. *p<0.05, **p<0.005, ***p<0.0005, ****p<0.00005.

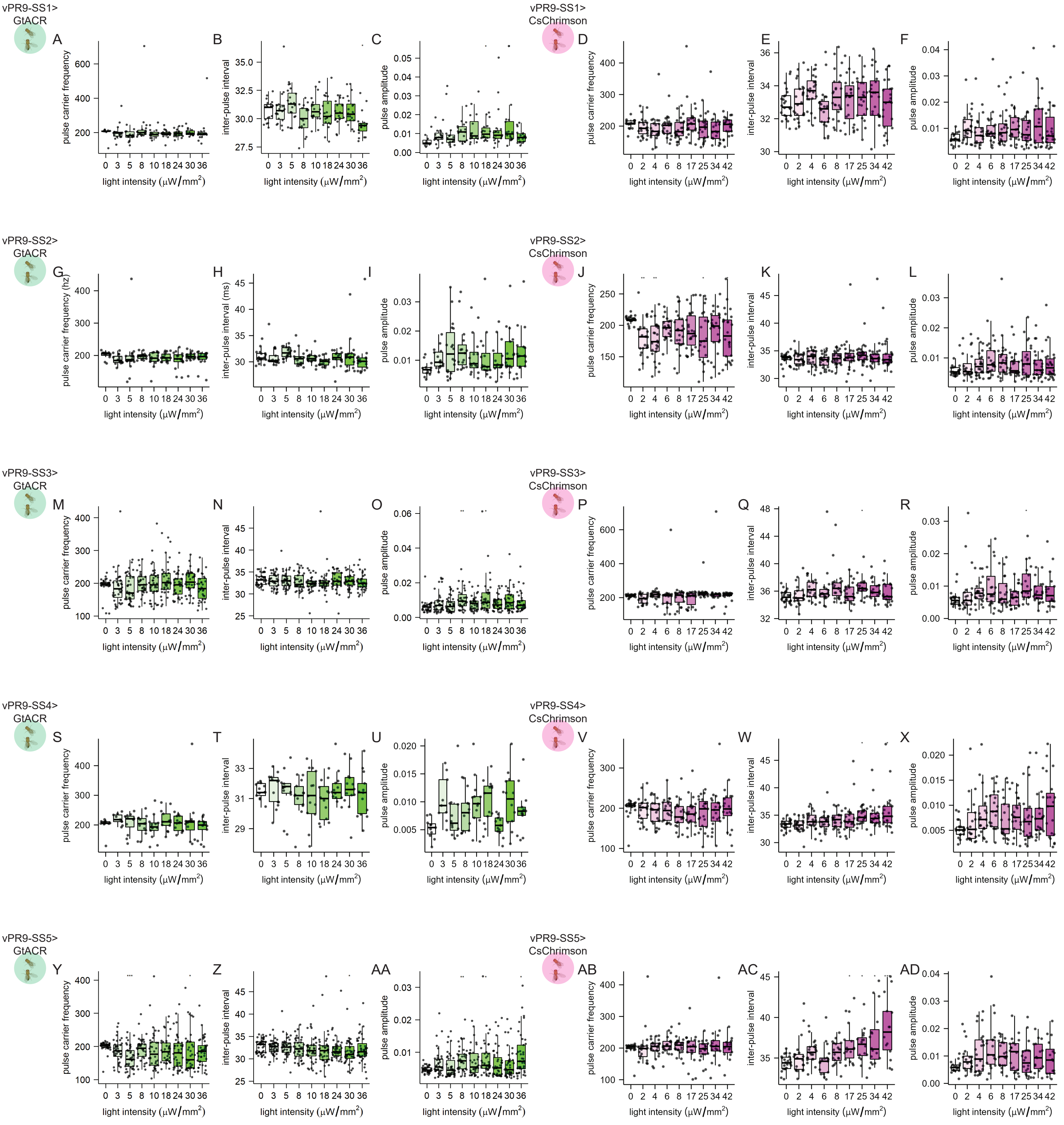


Sup. Figure 11 Quantification of pulse carrier frequency (hz), sine frequency (hz), pulse and sine amplitude (relative units), inter-pulse interval (ms), or polycyclic pulses/minute in courting split-GAL4>CsChrimson or split-GAL4>GtACR1 males without light stimuli (0) and increasing light stimulus intensities. split-GAL4 driver/neuron type, effector, and assay indicated to the left of each set of plots. p-values calculated via Kruskal-Wallis tests and Dunn's pairwise comparisons with Bonferroni correction. *p<0.05, **p<0.005, ***p<0.0005, ****p<0.00005.

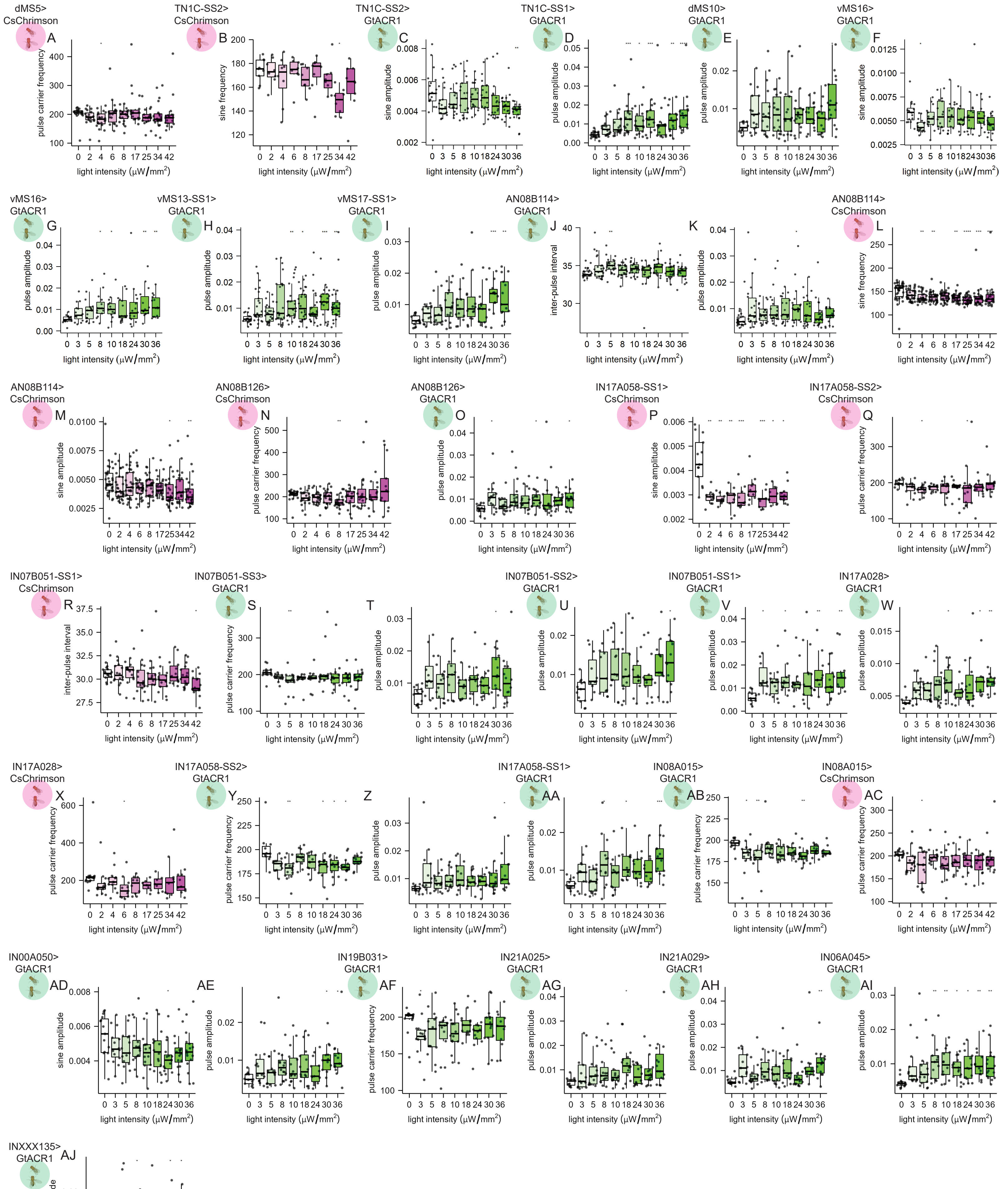

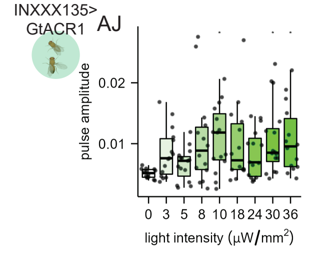


Sup. Figure 12 Quantification of pulse carrier frequency (hz), sine frequency (hz), pulse and sine amplitude (relative units), or inter-pulse interval (ms) in courting split-GAL4>CsChrimson or split-GAL4>GtACR1 males without light stimuli (0) and increasing light stimulus intensities. split-GAL4 driver/neuron type, effector, and assay indicated to the left of each set of plots. p-values calculated via Kruskal-Wallis tests and Dunn's pairwise comparisons with Bonferroni correction. *p<0.05, **p<0.005,***p<0.0005, ****p<0.00005.

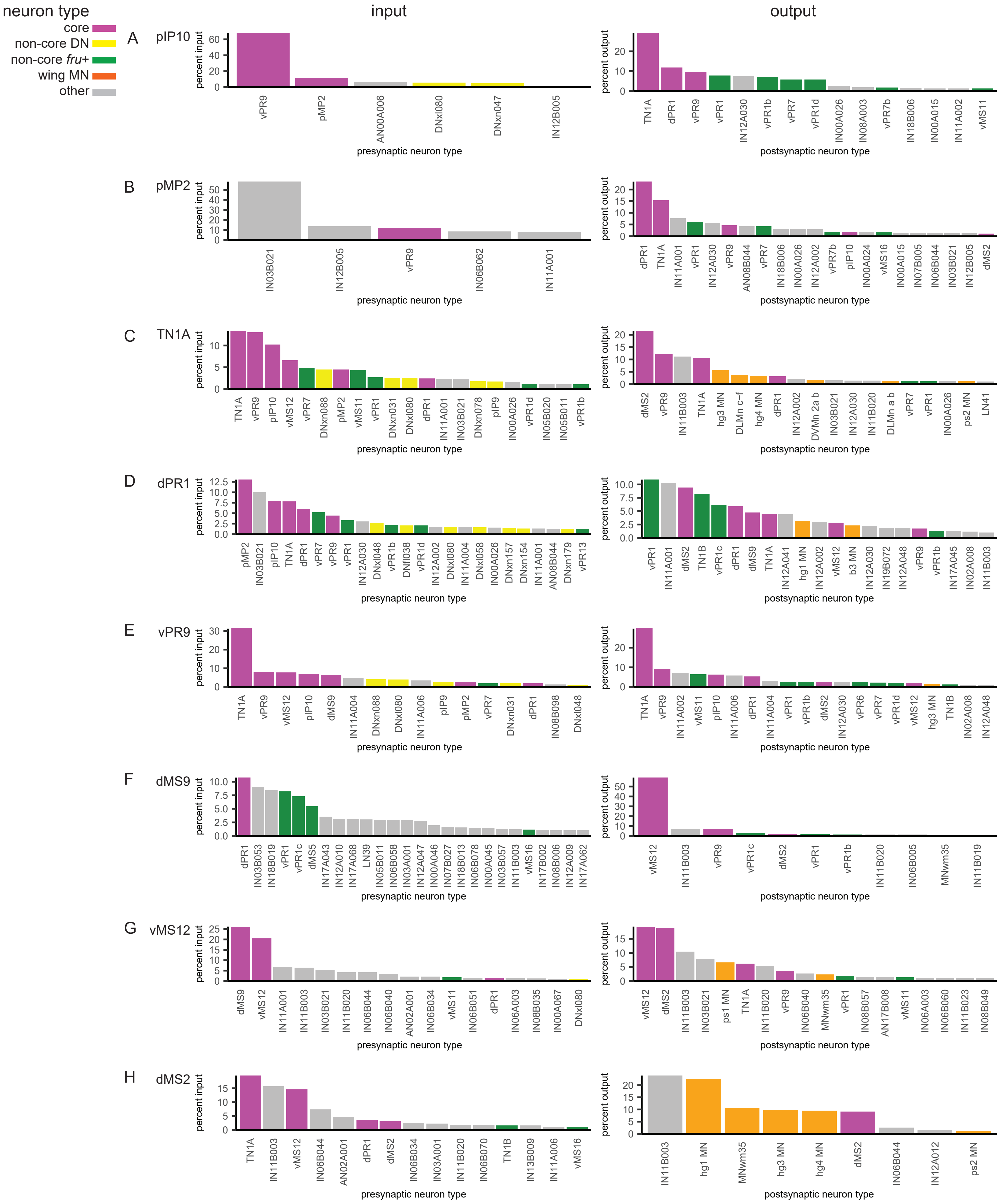


Sup. Figure 13: Connectivity to and from each core song circuit neuron. Every input and output of each core song circuit neuron with at least 10 connections was identified. These neurons were grouped by our manually assessed neuron type or systematic neuron type (MANC version 1.0). The percent of total input and output from each core neuron type was calculated. All neuron types with at least 1% input or output are shown.

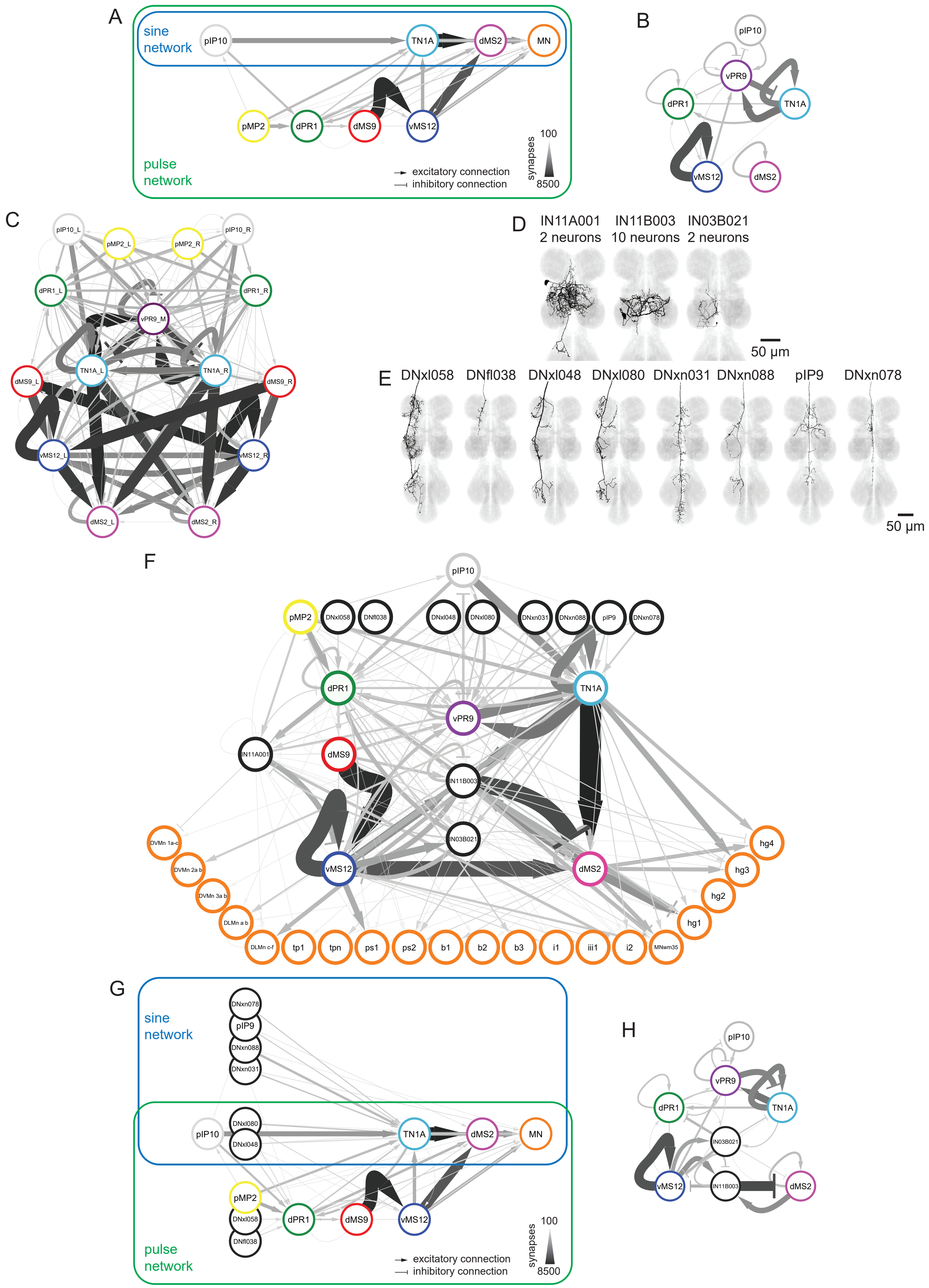
A
sine network
pulse network
pIP10
TN1A
dMS2
MN
pMP2
dPR1
dMS9
vMS12
excitatory connection
inhibitory connection
synapses
100
8500
B
vPR9
C
D
IN11A001 2 neurons
IN11B003 10 neurons
IN03B021 2 neurons
50 µm
E
DNxl058
DNfl038
DNxl048
DNxl080
DNxn031
DNxn088
pIP9
DNxn078
F
G
H
hg4
hg3
hg2
hg1
tp1
tpn
ps1
ps2
b1
b2
b3
i1
iii1
i2

Sup. Figure 14 A) Feed-forward pulse and sine networks within the core song circuit. vPR9 and within cell type connections were removed to make it easier to visualize the largely feed-forward connectivity. B) Recurrent connections within the core song circuit. pMP2, dMS9, and other purely feed-forward connections from (A) were removed to make it easier to view the recurrent connections. C) Core song circuit with connectivity between left right members of each cell type shown. D) Untested neuron types with >10% of the input to or output from at least two core circuit neurons. E) Untested descending neurons with relatively strong input to the core song circuit. F) The core song circuit, wing motor neurons, and neurons from (D) and (E). G) Same as (A), but with descending neurons from (E, white) that make connections to the pulse and sine feed-forward networks. H) Same as (B), but with neurons from (D, white) that make strong recurrent connections with core song circuit neurons.
